# Adaptive introgression and standing genetic variation, two facilitators of adaptation to high latitudes in European aspen (*Populus tremula* L.)

**DOI:** 10.1101/2021.02.23.432466

**Authors:** Martha Rendón-Anaya, Jonathan Wilson, Sæmundur Sveinsson, Aleksey Fedorkov, Joan Cottrell, Mark E.S. Bailey, Dainis Ruņģis, Christian Lexer, Stefan Jansson, Kathryn M. Robinson, Nathaniel R. Street, Pär K. Ingvarsson

## Abstract

Understanding local adaptation in plants from a genomic perspective has become a key research area given the ongoing climate challenge and the concomitant requirement to conserve genetic resources. Perennial plants, such as forest trees, are good models to study local adaptation given their wide geographic distribution, largely outcrossing mating systems and demographic histories. We evaluated signatures of local adaptation in European aspen (*Populus tremula*) across Europe by means of whole genome re-sequencing of a collection of 411 individual trees. We dissected admixture patterns between aspen lineages and observed a strong genomic mosaicism in Scandinavian trees, evidencing different colonization trajectories into the peninsula from Russia, Central and Western Europe. As a consequence of the secondary contacts between populations after the last glacial maximum (LGM), we detected an adaptive introgression event in a genome region of ∼500kb in chromosome 10, harboring a large-effect locus that has previously been shown to contribute to adaptation to the short growing seasons characteristic of northern Scandinavia. Demographic simulations and ancestry inference suggest an Eastern origin - probably Russian - of the adaptive Nordic allele which nowadays is present in a homozygous state at the north of Scandinavia. The strength of introgression and positive selection signatures in this region is a unique feature in the genome. Furthermore, we detected signals of balancing selection, shared across regional populations, that highlight the importance of standing variation as a primary source of alleles that facilitate local adaptation. Our results therefore emphasize the importance of migration-selection balance underlying the genetic architecture of key adaptive quantitative traits.

## Introduction

Local adaptation, the means by which populations of a species genetically adjust to local environments, is a powerful process of evolution. It occurs because multiple environmental factors imposing different selective pressures exist and the strength of each factor varies across habitats. When a population colonizes a new habitat, certain environmental conditions will impose higher selective pressures, while natural selection may be relaxed for other environmental factors. The overall shift in the selection landscape leads to local adaptation and consequent fitness trade-offs [1]. Two fundamental genetic sources for local adaptation, particularly in temperate forest trees that have large effective population size and low nucleotide substitution rates per unit of time, are standing variation and intra/inter-species hybridization. Hybridization occurs when reproductive isolation is not complete between species or when species lineages that are separated geographically meet after a secondary contact. Species capable of hybridization will therefore have access to a larger pool of genetic variation that provides the raw material for selection and accelerated adaptation. At the same time, standing variation can be maintained through balancing selection (BS) within populations. The signatures of BS include increased diversity around the target of selection, differentiation between populations departing from the genome-wide average and increased linkage disequilibrium, among others [2]. When selection varies geographically, it may favor locally adapted alleles in the derived lineages or subpopulations, giving rise to genomic regions with elevated F_ST_ and absolute divergence D_XY_ [3, 4].

Throughout the entire Pleistocene, range expansions and contractions of different species occurred in Europe, influencing the current patterns of diversity in many taxa [5]. Paleoclimate model simulations and pollen and plant macrofossil records have revealed a long-term decline in tree populations, with a threshold at the time of Heinrich Stadial (HS) 2 (24 thousand years ago [ka]), when tree populations decreased dramatically and did not recover until the end of HS1 (15ka). At this point, there were no refugia for temperate trees north of 45°N in Europe, which means that present day populations of temperate trees in northern Europe are essentially young, in contrast to conifers and other boreal tree species [6]. This also implies that temperate trees in northern Europe derive from southern populations, where they survived in glacial refugia until temperature and moisture conditions allowed for the recolonization of higher latitudes [7]. The distribution of glacial refugia across the continent and the successive contact of isolated populations following range expansions often led to hybridization events that played a major role in the evolution of forest trees and other plants [8, 9].

European aspen, *Populus tremula,* is a dioecious temperate angiosperm tree with a very extensive range across both the European and Asian continents. Its role in many ecosystems is that of a primary colonizer for forests, growing quickly and being able to cover large areas across both different latitudes and elevations [10]. Natural variation of *P. tremula* along a north-south gradient on the Scandinavian peninsula has been widely studied. GWAS and population differentiation analyses have identified a region on chromosome 10 harboring the *PtFT*2 locus, a gene known to be involved in controlling seasonal phenology [11], as a key contributor to climate adaptation at high latitudes in Sweden. Other loci harboring genes related to senescence processes or plant growth have also been proposed to have played a role in local adaptation, based on differences in haplotype frequencies identified with the hapFLK statistic [12] between Swedish populations originating at different latitudes [13].

Introgression has been hypothesized to be a major driver of adaptation in the genus *Populus*, where the lack of reproductive barriers allows for prevalent inter-species hybridizations. One such example is given by the adaptive introgression of a telomeric region on chromosome 15 from *P. balsamifera* to *P. trichocarpa* that has been implicated in climate adaptation in the recipient species [14]. In the case of *P. tremula*, admixture in contact zones along the Scandinavian peninsula has been documented and, even though different genetic lineages of *P. tremula* can be defined across Europe [15], no cases of adaptive introgression have been described so far in this species.

Using whole-genome re-sequencing data from an extensive set of Eurasian samples of *P. tremula* we describe different aspects of the post-glacial colonization, adaptation and admixture in the species across the Eurasian continent. We dissect the population structure of the species, revealing a complex and intricate genomic mosaicism supporting and extending earlier results from much more limited datasets [15]. Furthermore, we studied different types of selection that underlie local adaptation of aspen. Interestingly, we observed a clear and unique case of adaptive introgression, in which the strong selective sweep previously observed on chromosome 10 that harbors a gene controlling bud set in Northern Scandinavia resulted from a recent introgression event from the Russian gene pool. Overall, our results highlight the importance both of standing genetic variation and of genomic introgression as sources of new alleles for local adaptation.

## Materials and methods

### Sample collection and sequencing

The Swedish samples used in this paper belong to the SwAsp collection which has been extensively described previously [11, 16]. Leaf samples were collected from the common garden at Sävar [16] for all but two genotypes that were present only at the Ekebo common garden, for which wood tissue was sampled. Sampled tissue was stored in cool conditions until frozen at the laboratory. An additional 296 samples of *P. tremula* were obtained from across Eurasia. Trees selected from the U.K. were cloned from root cuttings and grown in one of two common gardens [17]: a Scottish national aspen collection maintained by Forest Research at Bush, Midlothian, and the Eadha Enterprises clone garden at Lochwinnoch, Renfrewshire. Leaves were sampled from 125 genotypes at Bush and 15 genotypes at Lochwinnoch, representing a range of national seed zones defined by the UK Forestry Commission. Samples were freeze-dried prior to transportation to Sweden. In Norway and in Russia, leaf material was sampled from 24 trees within a region. In Latvia, ten trees were sampled in each of five regions spanning the country. Fewer samples were obtained from Iceland due to the limited availability of natural stands of *P. tremula*. Due to the clonal growth of *P. tremula*, individual samples were collected with enough physical separation (0.3 km) between individuals to minimize the risk of sampling identical genets. Leaf samples were stored in silica gel before shipping to Sweden. Geographical coordinates were recorded for the sampled trees. The individuals included in the study (after the quality control steps that follow) and their geographic origins are listed (Additional File2: Table S1).

Total genomic DNA was extracted from frozen leaf tissue for all individuals using the DNeasy plant mini prep kit (QIAGEN, Valencia, CA,USA). For the Icelandic samples, library preparation and sequencing were conducted by Génome Québec, Canada. Briefly,1 µg of high-quality DNA was used for paired-end libraries construction. The 12 libraries were sequenced in three lanes of Illumina HiSeqv4 with read length of 125 bp. All other samples were subjected to paired-end sequencing using libraries with an average insert size of either 350bp or 650bp and samples were sequenced by the National Genomics Infrastructure at Science for Life Laboratory, Stockholm, on an Illumina HiSeq 2000 or Illumina HiSeq X platform to a mean per-sample depth of approximately 17X.

### Mapping and SNP calling

Re-sequenced accessions were mapped against the reference genome of *P. tremula* v2.0 [18], using BWA (v0.7.17; [19] – mem alignments for paired-end reads using default parameters. Post-mapping filtering removed reads with MQ<20 (using samtools v1.10; [20]. Depth and breadth of coverage were assessed in order to confirm all samples had a minimum coverage of 8X (Additional File2: Table S1). Finally, before the variant calling step, we tagged duplicate reads (using Picard MarkDuplicates v2.10.3; Picard Toolkit.” 2019. Broad Institute, GitHub Repository: http://broadinstitute.github.io/picard/) and found that they did not exceed 14% of the sequencing reads in individual libraries (ranging from 3 to 13.8%). We used GATK v3.8 to call variants [21]. First, we performed a local realignment around indels with RealignerTargetCreator and IndelRealigner (using default parameters). We called per-sample variants using HaplotypeCaller to produce gVCF files (-ERC GVCF). Given the large number of individuals, we produced intermediate gVCF files using CombineGVCFs that were finally used in the joint-call step with GenotypeGVCFs. From the resulting VCF file, we selected SNPs using SelectVariants and filtered them with VariantFiltration (QD < 2.0; FS > 60.0; MQ < 40.0; ReadPosRankSum < -8.0; SOR > 3.0; MQRankSum < -12.5). At this point, the filtered VCF was lifted over to version 2.2 of the genome of *P. tremula*, available at ftp://plantgenie.org/Data/PopGenIE/Populus_tremula/v2.2/, using picard LiftoverVcf, which allowed for better resolution at the chromosome level. Further SNP pruning with vcf/bcftools removed positions with extreme depth values (min-meanDP 10, max-meanDP 25; these thresholds correspond to the average depth ± one standard deviation), absent in more that 30% of the samples, non-biallelic or displaying an excess of heterozygosity (FDR <0.01).

### Batch removal on SwAsp individuals

Given the fact that two sequencing batches from different Illumina platforms (Illumina HiSeq 2000 and HiSeq X) were used to cover the SwAsp collection, we observed noisy signals at the population structure level, presumably as a result of the different sequencing equipment and library preparation methods. By means of principal component analyses (PCA) we identified differences in the grouping of the individuals where differences among samples along PC1 and, to a lesser extent, PC2 were explained by the sequencing platform and not the geographic origin of samples (Additional File1: Figure S1A; PC1 separates the SwAsp collection in two sets, each corresponding to a different Illumina sequencing platform). To address these batch effects, we removed SNPs associated with both components without losing the geographic structure of the Swedish population. For this purpose, we generated a genotype file with vcftools for each chromosome. We ran independent PCA analyses on each SNP configuration (homozygous reference, heterozygous, homozygous alternative) and realized it was at the heterozygous level where the batch effect was present. Using the libraries “ggfortify”, “factoextra” and “FactoMineR”, we estimated the contribution of each variant to the components affected by the platform effect. We started by assuming a uniform contribution of each variant to each component, and removed those that would deviate from this premise in both PC1 and PC2 using the following rational:

Assuming uniformity in the contribution of the variants, we calculate the expected C value:

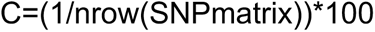

Threshold (T) of variants contribution both to PC1 and PC2 under uniformity:

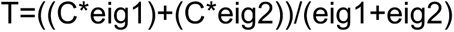

(where eig1 and eig2 are the eigenvalues corresponding to PC1 and PC2)

For each variant, we calculate the contribution value:

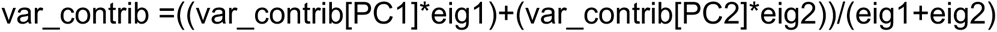

We kept only those variants that did not contribute more than expected under the uniformity hypothesis (var_contrib <= T). While removing the batch effect using this procedure, we observed loss of more than 4e^6^ SNPs (Additional File1: Figure S1B).

In order to evaluate if we could remove the batch effect without compromising such a large number of variants, we assigned a p-value to the contribution of each SNP to the components using dimdesc (FactoMineR) at the chromosome level. We tested different cut-offs (Additional File1: Figure S2) and observed that the batch effect was removed while maintaining the geographic distribution of samples when using a p-value threshold of 0.05 for PC1 and 0.01 for PC2 (Additional File1: Figure S2B). With these thresholds, ∼1.8 e^6^ SNPs were filtered out to remove the batch effects but without compromising the overall population structure of the samples.

### Population structure and admixture in *P. tremula*

We used plink (v1.90b4.9; [22]) to identify linked and low frequency SNPs (--indep-pairwise 100 10 0.2 –maf 0.05) that we removed with vcftools. We used the resulting set of pruned positions to compute the relatedness between samples from the Eurasian collection by calculating genome-wide estimates of identity by descent (IBD). We removed from our downstream analyses one sample from each pair of closely related individuals, using a threshold value of relatedness of 0.4.

Next, we used vcftools to output the genotype likelihood information contained in the pruned VCF file; the resulting beagle-formatted file was input into NGSAdmix to estimate individual admixture proportions across a varying number of ancestral populations (K=3 to K=6).

Finally, we assessed individual ancestries using EIGMIX implemented in SNPRelate. This eigen analysis, developed by [23], is computationally efficient for estimating ancestral proportions by making assumptions of surrogate samples for ancestral populations. For this, we chose the Latvian, Scottish and Russian populations as proxies of the ancestral populations (referred to as Central European, Western European and Siberian, respectively) from which admixture was to be estimated, based on earlier estimates of the post-glacial colonization history of *P. tremula* across Europe [15]. We used the LD/MAF pruned SNPs and produced a GDS object for the analysis [24].

### Gene flow

We combined several statistics developed to identify introgressed genomic regions. These included the classic ABBA-BABA test in its developed F_d_ statistic form. For these calculations we used the popgen pipeline developed by Simon Martin, available at https://github.com/simonhmartin/genomics_general. We treated each chromosome independently; the corresponding vcf file was converted to .geno format using parseVCF.py. From these files, we calculated diversity and divergence values for each subpopulation (D_XY_, F_ST_ and π) in non-overlapping 10 Kb windows, using the script genomics.py (Additional File2: Table S2). Finally, we used the script ABBABABAwindows.py to compute the four-taxon D statistic and f estimators in non-overlapping windows of 10 Kb as well. For these calculations, we tested gene flow from the Russian subpopulation (P3; 23 individuals) into Northern Scandinavia (P2; 56 individuals: 34 from Northern Sweden and 22 from Norway), setting the Chinese (O; 15 individuals) samples as the outgroup, and the Latvian (45 individuals) or Southern Scandinavian (50 from Sweden and 23 from Norway) groups as the other potential receptors (P1). Since we were interested in estimating shared variation between Russian and Northern Scandinavian individuals, we focused on the f_dM_ introgression statistic, described by [25]. f_dM_ gives positive values when introgression occurred between P3 and P2, and negative values if it occurred between P3 and P1. In order to avoid stochastic errors that could produce meaningless values, we only considered windows with at least 100 biallelic SNPs.

We also calculated the basic distance fraction, Bd_f_ (PopGenome; v2.7.1; [26], which combines both f_d_ and distances estimations and that is less sensitive to the time of gene-flow. Just as in the previous analysis, we set a four-taxon tree hypothesis to estimate the proportion of introgression from Russia into Northern Scandinavia. We calculated Bd_f_ in 10 Kb windows treating each chromosome independently. We combined all 19 chromosome estimations and assigned a Z-score and p-value to each Bd_f_ value using genome-wide results and did an extra FDR correction. We selected those regions that had an FDR<0.05.

Finally, we ran the Efficient local ancestry inference (ELAI) method [27] to confirm the introgression event on chromosome 10. We used two upper-layer clusters and 10 lower-level clusters; 20 Expectation-Maximization steps and 100 generations of admixture between the ancestral populations, that corresponded to the Russian and Latvian aspen populations. The plotted allele dosages correspond to the average over all the Swedish individuals from the northern population. We ran five independent EM runs.

### Positive and balancing selection

We scanned the genome of *P. tremula* for signals of positive selection using a newly developed strategy called “integrated selection of alleles favoured by evolution”, iSAFE [28]. This method first calculates haplotype allele frequency scores based on the presence of derived alleles in a particular haplotype, which is then used to calculate SAFE scores. These SAFE scores are in turn calculated across a region of given size in 50% overlapping windows of 300 bp to culminate in an iSAFE signal. These statistics can be calculated for large regions of phased haplotypes, which we obtained with BEAGLE v.4.1 [29], so chromosomes were divided into 3 Mb windows for each iSAFE iteration. The iSAFE software can be set to run under a case-control mode, with the case populations being, for example, the Northern Scandinavian population, and the control population being all remaining individuals not used in the case population. We ran this screening for Northern and Southern Scandinavia, Russia, and combining all the Nordic individuals from Norway, Sweden and Russia together. As recommended by the authors, we considered iSAFE values significant when they were >0.1.

In addition, we ran betascan [30],to detect possible signals of balancing selection, by dividing the collection in geographic zones: Northern Scandinavia (Sweden and Norway), Central Europe (Latvia and Southern Sweden), Western Europe (Scotland) and Southwestern Scandinavia (Southern Norway and Iceland), and using the following parameters: -fold -m 0.1 -p 20. To avoid any bias due to the unbalanced number of individuals in each group, we randomly chose 55 individuals from each geographic cluster. In order to avoid spurious signals of balancing selection, we binned SNPs with high beta-scores (FDR<0.01) into 50 Kb windows and searched for genes encoded within bins having at least 100 significant SNPs. Functional enrichments were obtained using topGO v.2.36.0 [31].

### Population demographics

We chose StairwayPlot [32] to infer changes in effective population sizes in the past. For this, the folded site frequency spectrum (SFS) for the tested subpopulation was calculated using ANGSD v.0.920 [33] using the LD-pruned SNPs (plink --indep-pairwise 100 10 0.2). We ran independent analyses for Scandinavia, Russia, Latvia and for a set of 50 randomly selected individuals from the entire collection. We generated 100 bootstrap replicates and assumed a mutation rate (μ) of 2.5e-9 and a generation time of 15 years [34].

In addition, we aimed to confirm the directionality of the gene flow events we detected on chromosome 10 between the Russian and Scandinavian individuals. For this, we explored alternative demographic models using the diffusion approximation method of dadi [35] to analyze the site frequency spectra of our aspen subpopulations. We used two chromosomes for this purpose, chromosome 10 and chromosome 8, since there is strong evidence of synteny between both chromosomes deriving from the 40 My old genome duplication that are shared by all member of the genus *Populus* [36]. We ran analyses independently for chromosome 8 and 10 using all LD pruned, biallelic sites including all biallelic sites present in the putatively selective sweep/introgressed region on chromosome 10. We fit 19 demographic models (Additional File1: Figure S3) for the Russian and Scandinavian subpopulations using the demographic modelling workflow (dadi_pipeline) from [37]. We tested different classes of models: simple models of divergence with and without migration; simple models plus instantaneous size changes, ancient migration or secondary contact, ancient migration plus instantaneous size change, and island models of vicariance and founder events. In total, we tested 19 different demographic scenarios using all polymorphic sites on chromosome 10. First, we ran the general optimization routine (dadi_Run_Optimizations.py), which includes fitting the model using particular settings for a given number of replicates, then using the parameters from the best scoring replicate to seed a subsequent round of model fitting using updated settings. We used four rounds, with 10, 20, 30 and 40 replicates. The arguments controlling the steps of the optimization algorithm and the perturbation of starting parameters were maxiter=[3,5,10,15] and folds=[3,2,2,1]; we defined the extrapolation grid size to pts = [80,90,100] and projection sizes to proj = [15,15]. The optimization routine with four rounds had an important effect in minimizing differences in the likelihoods achieved at the end of the runs (Additional File1: Figure S4). In order to generate confidence intervals for the parameters estimated from the demographic models with the highest likelihood, we created 100 bootstrap replicates of the spectra and calculated the standard deviations of the best-fit parameters using the Godambe methods [38].

## Results

### Population structure and admixture

In order to elucidate demographic and adaptive events that have accompanied the post-glacial radiation of European aspen, we re-sequenced a large collection of individuals, covering a wide geographic range across the Eurasian continent. Our collection comprises 411 *P. tremula* individuals sampled from China [39], Russia, Latvia, Norway, Sweden, Iceland and Scotland (Figure 1A; Additional File2: Table S1). Sequencing reads for all individuals were mapped against the reference *P. tremula* v2.0 genome [18] and obtained ∼20.8e^6^ SNPs after several filtering steps (depth of coverage, missingness, level of heterozygosity and batch effect caused by the incorporation of samples in the Swedish collection from two different sequencing platforms; Additional File1: Figure S1,2). It should be highlighted that the removal of false heterozygous positions did not alter the results of the downstream analyses (Additional File1: Figure S5). Finally, we removed 48 samples that were possible hybrids or that were highly related samples, yielding a total of 363 individuals for all downstream analyses.

**Figure 1.**
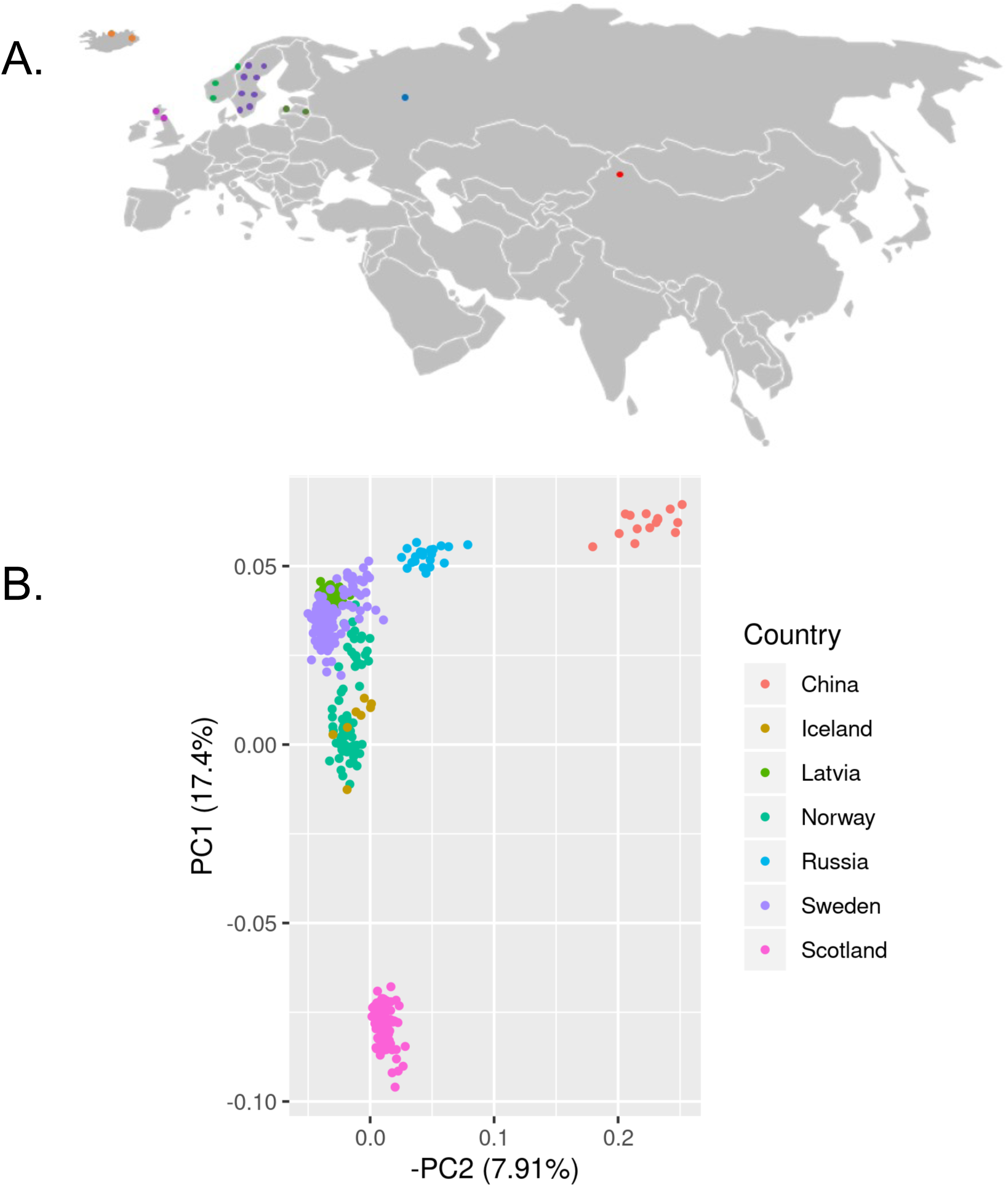
*P. tremula* collection. A. Sampling sites across the Eurasian continent. B. PCA of Principal component analysis of *P. tremula* individuals, using 283,505 pruned sites across the 19 chromosomes.

Due to the broad geographic coverage of *P. tremula*, we first focused on estimating the population structure of the species. As depicted in Fig 1B, a principal component analysis (PCA) using 2.8e^5^ pruned, unlinked SNPs clearly separated at least three independent clusters of individuals, one corresponding to samples of Chinese origin, another comprising all samples collected across continental European and a third comprising samples collected on the British Isles (primarily Scotland). A deeper analysis of the composition of the species structure with NGSAdmix revealed five different ancestral populations and different levels of hybridization in particular geographic regions. Chinese and Russian samples, represented by solid colors in Figure 2A (orange and yellow, respectively), did not show strong signs of admixture with European samples. However, when moving towards Western Europe, admixture events occur and we can recognize three additional populations: a Scandinavian (purple), a Central-European (Latvia, blue) and a Western European (Scottish, green) population. Interestingly, the genomic background of the Scandinavian samples (from Norway and Sweden) is a complex mosaic of up to four populations, confirming earlier results based on microsatellite data [15] and providing additional evidence for several waves of migration that led to the colonization of the Scandinavian peninsula.

**Figure 2.**
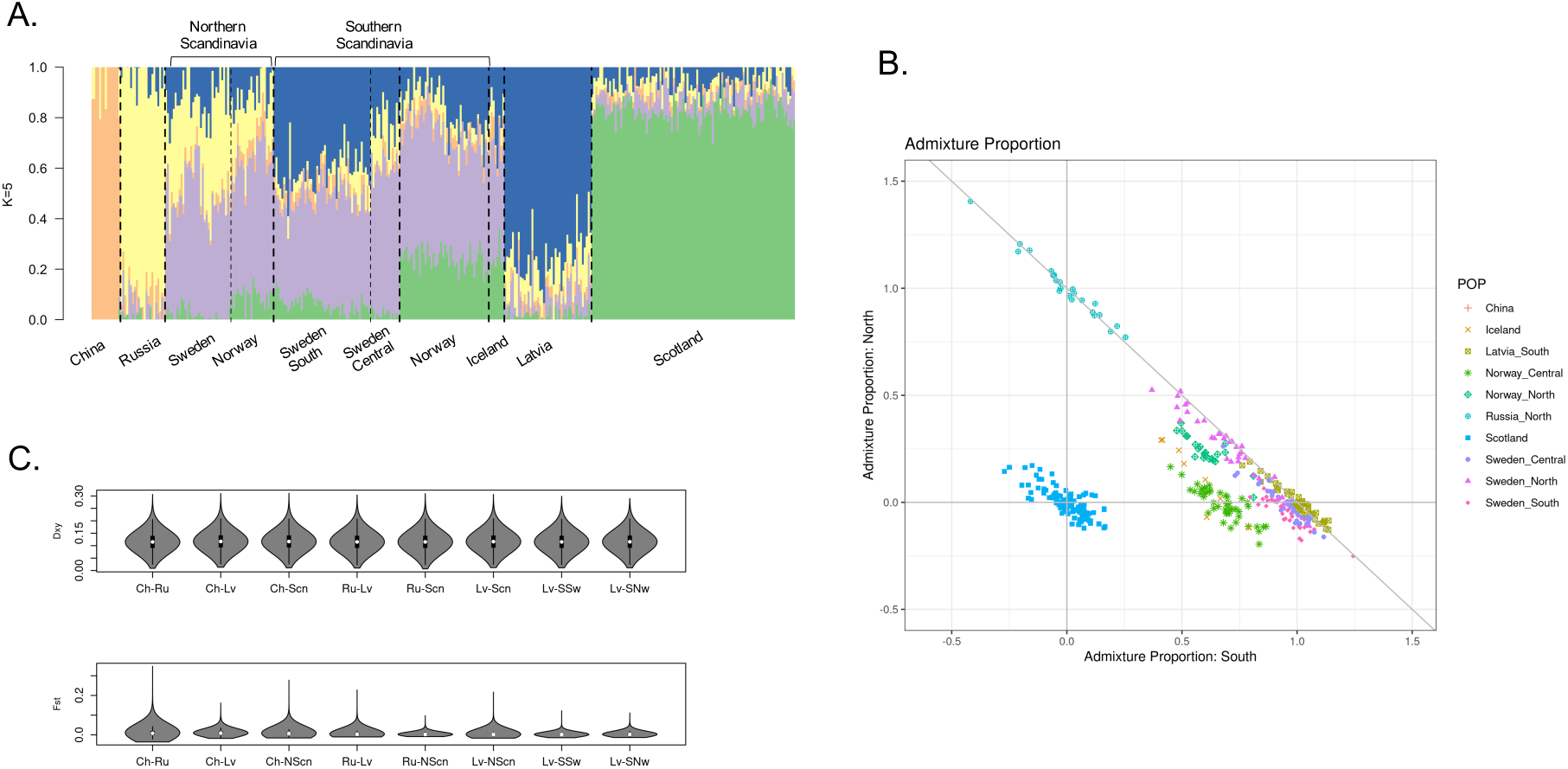
Population structure and admixture proportion. A. NGSadmix plot (K=5) showing the hybrid genetic background of the European populations. B. Admixture plot (EIGMIX) where the Russian, Scottish and Latvian populations where selected as proxies of the ancestral gene-pools in Europe and thus, are located at the vertices of the plot. C. Diversity and differentiation estimators between aspen subpopulations.

Given these observations, we took a closer look at the genomic mosaicism of the Scandinavian population. We used EIGMIX, taking Russian, Scottish and Latvian populations as proxies of the ancestral Siberian, Western and Central European populations, respectively (Figure 2B, see also [15]). These populations correspond to the vertices of the graph in Figure 2B, while the remaining samples were positioned according to their level of admixture along the sides of the triangle. This analysis, consistent with the observations derived from the NGSadmix analysis, shows that individuals collected in Northern Scandinavia have on average 26% of Russian origin with a maximum value of 52% in Sweden (SWASP_108 and SWASP_115) and 37% in Norway (NO_MIR20). While the second highest ancestral component (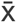 =0.66) in Northern Scandinavia corresponds to the Central European population, in the samples from northern Norway there is up to 20% of ancestry from the Western European population. The opposite trend was observed in Southern Scandinavia, where the dominant ancestral component is Central European (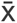 =0.75), with less than 10% of Russian admixture and up to 18% and 44% of Western European ancestry in samples from Sweden and Norway, respectively. This Western European genomic component (colored in green in Figure 2A), which is strongly represented among the Scottish and Norwegian samples, derives from an additional ancestral population, probably located between the Iberian and Italian peninsulas that we did not cover in our sampling. These observations also fit the results from earlier analyses based on microsatellite data [15]. Icelandic samples largely overlap with Norwegian individuals, suggesting that the population in Iceland has been recently introduced and has a clear Norwegian origin.

These observations were also supported by F_ST_ estimations (Figure 2C, Table 1), as we obtained a small dispersion of values when calculating pairwise comparisons between Russia and Northern Scandinavia (mean F_ST_=0.003, maximum at 0.09), or Latvia and Southern Scandinavia (mean F_ST_ between Latvia and Sweden of 0.002, maximum at 0.07 and between Latvia and Norway of 0.004, maximum at 0.1), while comparisons between populations without signs of admixture, such as China and Russia, reached values as high as 0.35, which approach F_ST_ values observed between closely related aspen species [39].

**Table 1.** Genome-wide F_ST_ comparison.

|  |  | CH | ICE | LV | NOC | NON | NOS | RU | SWC | SWN | SWS | UK |
| --- | --- | --- | --- | --- | --- | --- | --- | --- | --- | --- | --- | --- |
| $F_{ST}$ | CH | - | | | | | | | | | | |
|  | ICE | 0,014 | - |  |  |  |  |  |  |  |  |  |
|  | LV | 0,012 | 0,003 | - |  |  |  |  |  |  |  |  |
|  | NOC | 0,016 | 0,003 | 0,005 | - |  |  |  |  |  |  |  |
|  | NON | 0,013 | 0,002 | 0,004 | 0,003 | - |  |  |  |  |  |  |
|  | NOS | 0,016 | 0,003 | 0,004 | 0,003 | 0,003 | - |  |  |  |  |  |
|  | RU | 0,012 | 0,006 | 0,005 | 0,007 | 0,005 | 0,007 | - |  |  |  |  |
|  | SWC | 0,016 | 0,004 | 0,003 | 0,004 | 0,003 | 0,003 | 0,006 | - |  |  |  |
|  | SWN | 0,012 | 0,003 | 0,004 | 0,004 | 0,002 | 0,003 | 0,004 | 0,001 | - |  |  |
|  | SWS | 0,011 | 0,002 | 0,002 | 0,003 | 0,002 | 0,002 | 0,005 | 0,001 | 0,002 | - |  |
|  | UK | 0,010 | 0,002 | 0,009 | 0,004 | 0,005 | 0,004 | 0,008 | 0,004 | 0,007 | 0,008 | - |

### Gene flow

We scanned the genome of *P. tremula* for potential introgression events that could have occurred given the admixed background we observed in Scandinavia. We followed a four-taxon approach for these scans, keeping the Chinese population as an outgroup in all our comparisons. We calculated the f_dM_ introgression statistic [25], and the basic distance fraction, Bd_f_ (PopGenome; v2.7.1), which combines both f_d_ and distance estimations and that is less sensitive to the timing of gene-flow. Both estimators give positive values when introgression occurs between P3 (donor population: Russia or Latvia) and P2 (Northern Scandinavia), and negative values if it occurred between P3 and P1 (Southern Scandinavia), on a scale from -1 to 1. When evaluating gene flow from Russia into Scandinavia, we obtained f_dM_ and Bd_f_ mean genomic values of -0.008 and 0.014, respectively (Figure 3A). After FDR correction at a threshold of 0.01, one region on chromosome 10, spanning from 4.5 to 4.9 Mb had a highly significant introgression signal, reaching values of f_dM_ =0.85 (FDR=3.6e^-3^) and Bd_f_ =0.89 (FDR=1.33e^-50^). As expected, there were differences in D_XY_ and F_ST_ patterns between subpopulations at this region: F_ST_ is remarkably high between Nordic subpopulations and any other geographic location, while D_XY_ decreases when northern Norwegian, Swedish and Russian subpopulations are contrasted (Additional File1: Figure S6,7). Introgression on chromosome 10 was further validated applying ELAI on each individual from the north of Sweden. This method implements a two-layer HMM (hidden Markov model) to infer local ancestry of admixed individuals without prior definition of window sizes or haplotype phasing, returning the most likely proportions of ancestry at each variable position of the chromosome. ELAI confirmed that, while chromosome 10 shares alleles from both ancestral or training populations and on average, northern individuals have 44% and 56% admixture proportions from Latvian and Russian populations respectively, the region encompassing 4.5-4.9 Mb has a predominant Russian ancestry (Figure 3B; Additional file 1: Figure S8).

**Figure 3.**
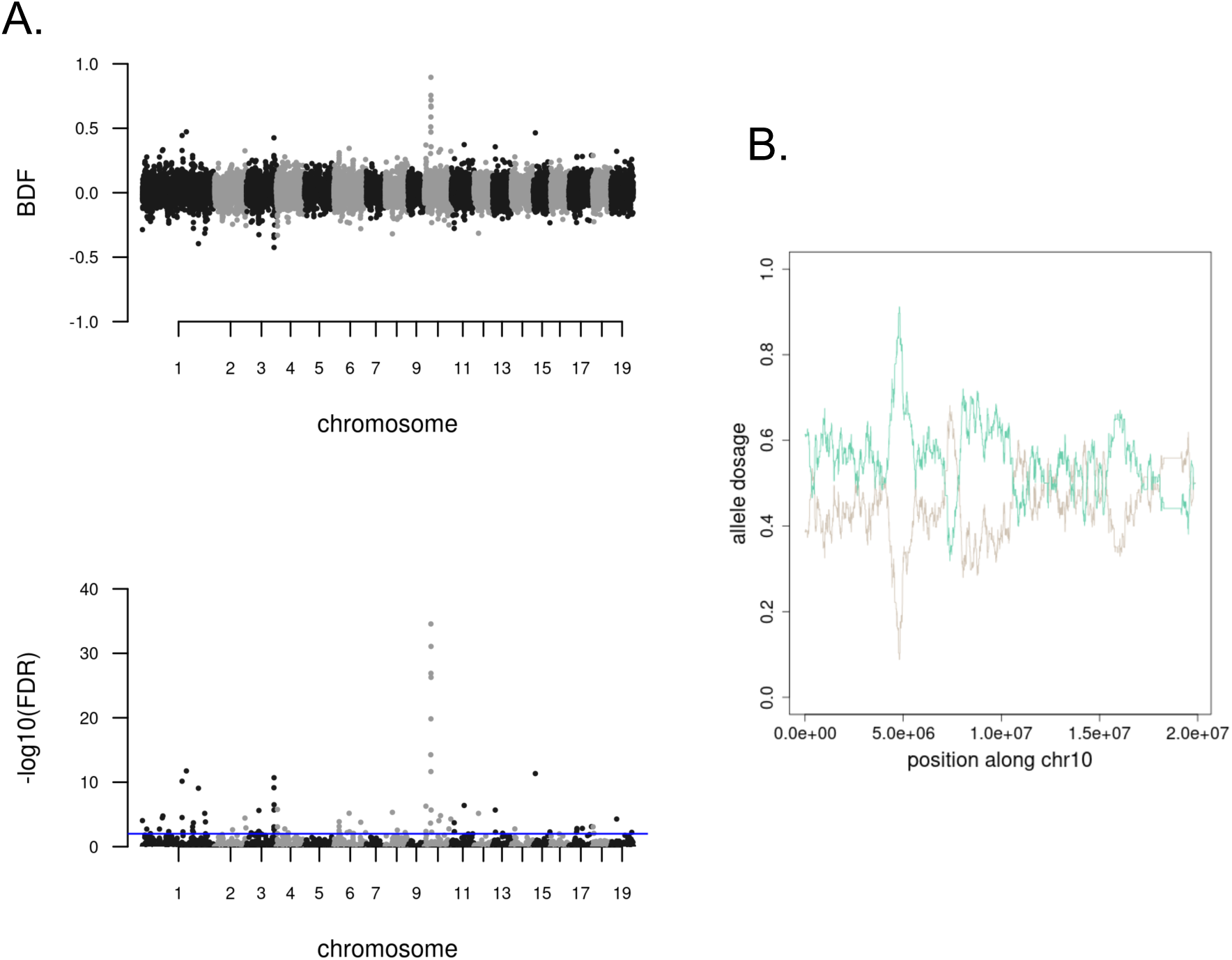
Introgression signals between Scandinavia and its ancestral gene pools. A. Gene flow from Russia using a four-taxon tree hypothesis. B. Allele dosage along chromosome 10 estimated with ELAI. In green, the dosage from the ancestral Russian population; in brown from the Latvian population.

In terms of gene content, two relevant loci annotated as Heading-date 3A-like, or Flowering Locus T (PtFT2) genes, are encoded in this introgressed genomic block: Potra2n10c20839 (4772117 - 4776457 bp) and Potra2n10c20842 (4789807 – 4792846 bp) and the region surrounding these two genes is known to have been subject to a selective sweep in the northern Swedish population [11].

### Alleles favored by evolution or positive selection

A good way to understand local adaptation without necessarily having phenotypic data is to look for signs of positive selection in different populations across a species range. Previous evidence of selective sweeps has shown a particularly strong signal on chromosome 10, in a region encompassing the two Flowering Locus T homologs [11]. This region is strongly implicated in the capacity of Nordic populations to adapt their seasonal phenology to the more variable day lengths and shorter growth seasons experienced at norther latitudes. We scanned the genome of *P. tremula* using a novel method to detect mutations favored by selection, iSAFE (integrated selection of alleles favored by evolution; [28]. We calculated iSAFE scores for each European population, and also contrasted Northern vs. Southern populations in Scandinavia in a case/control analysis mode. The highest iSAFE signal (iSAFE=0.22) was located on chromosome 10, in the region spanning 4.5 to 4.9 Mb where iSAFE values reached the recommended threshold of 0.1. The high iSAFE values were reached not only in Swedish individuals, but also when we combined samples originating in Northern Norway, Northern Sweden and Russia (Figure 4), implying that the sweep is shared among these populations derived from more northern latitudes in the Eurasian continent. As mentioned before, this region is centered around two loci annotated as Heading-date 3A-like, or *PtFT*2, in agreement with previous analyses that have associated *PtFT*2 with climate adaptation *in P. tremula* [11].

**Figure 4.**
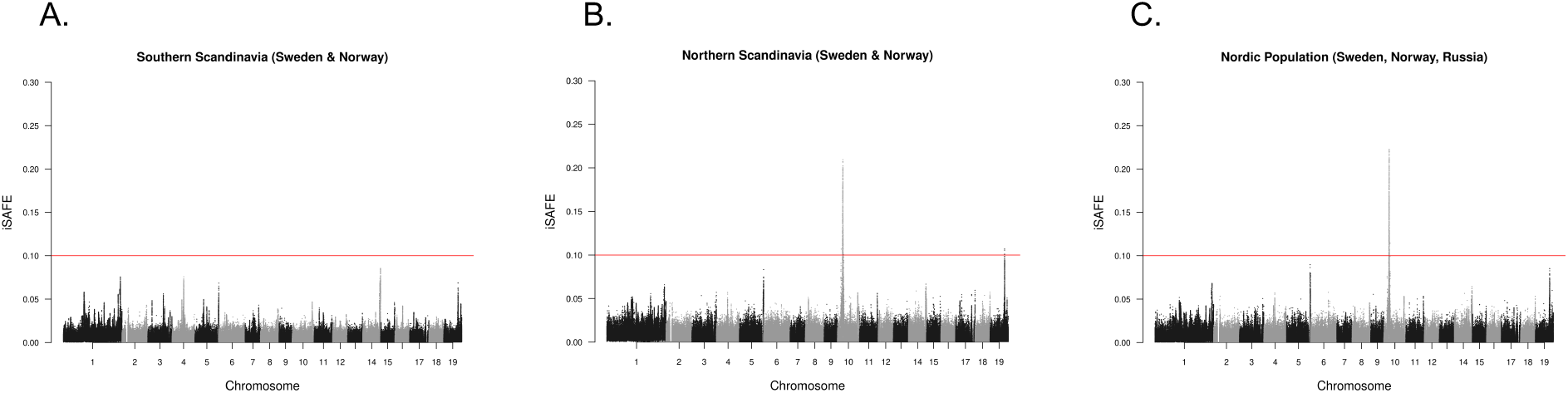
Selective sweep in chromosome 10 in Nordic populations. A. iSAFE scan of individuals from the southern part of Scandinavia. B. iSAFE scan of individuals from the northern part of Scandinavia. C. iSAFE scan of chromosome 10 grouping Swedish, Norwegian and Russian individuals.

### Balancing selection and functional enrichments

A closer look at the iSAFE patterns in the different subpopulations revealed several strong signals that a) did not reach the accepted significance threshold to be confidently classified as regions targeted by positive selection or b) were not geographically limited, i.e., they are present in at least two distant subpopulations (Additional File1: Figure S9). This suggests that another selective force may be acting to maintain alleles at high frequencies, for instance balancing selection (BS). In order to evaluate this possibility, we ran betascan [30] on all 19 chromosomes, dividing the collection in geographic zones: Northern Scandinavia (Northern Norway and Sweden), Central Europe (Latvia and Southern Sweden), Western Europe (Scotland) and Southwestern Scandinavia (Southern Norway and Iceland). We hypothesized that some genomic tracts with a high beta-score would match regions that we previously classified as iSAFE candidates for positive selection across multiple populations. This pattern was evident on chromosomes 1 (23.4-23.5 and 38.2-38.35 Mb), 4 (9.9-10Mb), 5 (8.35-8.4 Mb), 7 (9.1-9.15 Mb),14 (14,7-14.8 Mb),17 (8.85-8.9 Mb), 18 (9.4-9.45 Mb) and 19 (5.6-5.65 Mb) (Figure 5). A further validation of the effect of BS on these regions comes from the comparison of π and D_XY_ patterns. While the average of genome wide, pairwise D_XY_ was around 0.12, the average value obtained when considering only the chromosome tracts affected by BS was close to 0.14 (t-test, *p*<0.01; Additional File2: Table S3), while π values also increased in BS targeted regions compared to genome-wide average (Additional File2: Table S3). Average F_ST_ values ded not differ between BS affected regions and genome wide estimations, however.

**Figure 5.**
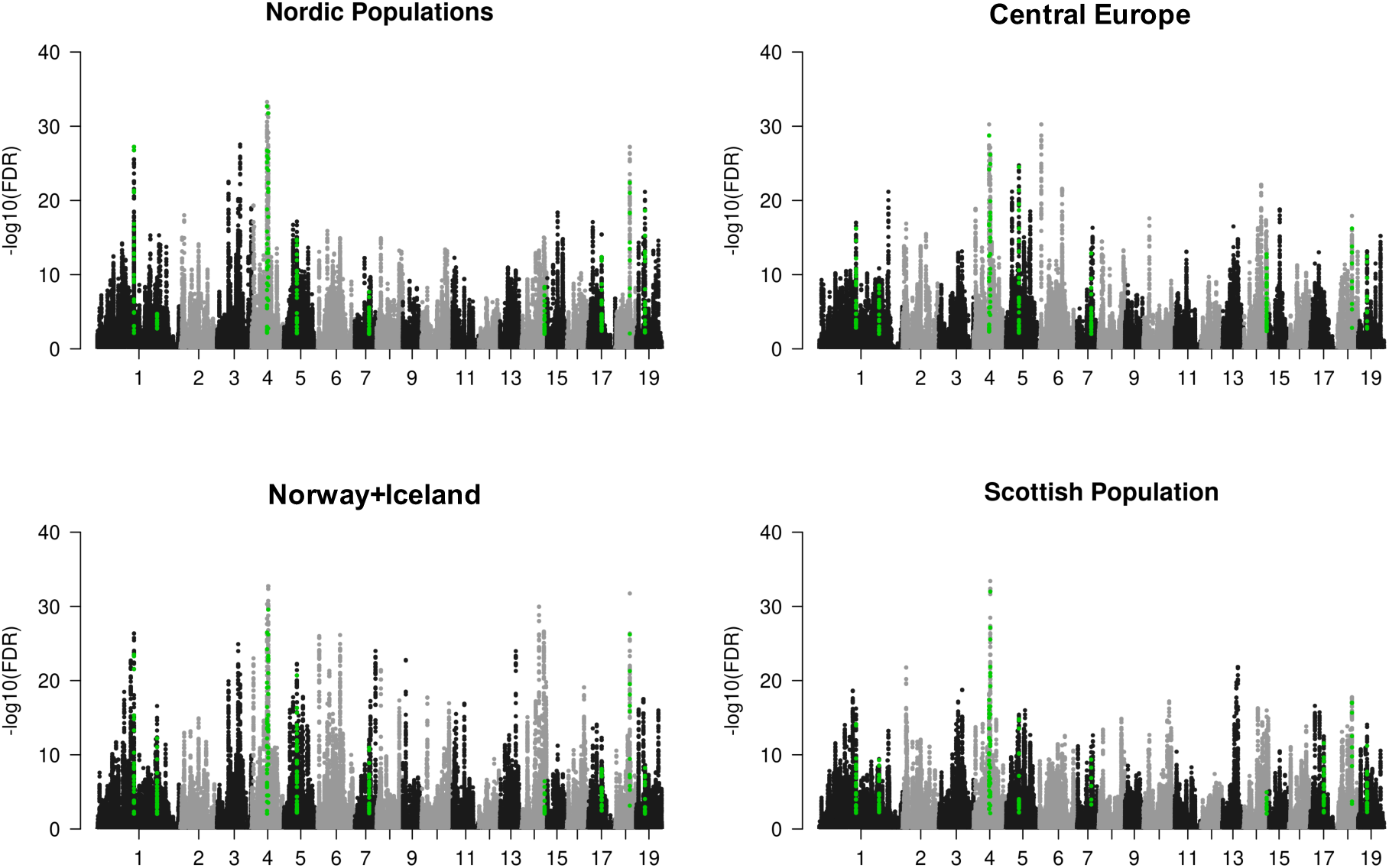
Balancing selection. Green dots highlight shared signals between subpopulations.

Furthermore, we also observed significant bins that were unique to a specific population and we therefore ran independent functional enrichment tests for biological processes affected by balancing selection for each of these regions. Interestingly, we found a few common terms with significant p-values (Fisher test, *p*<0.05), such as ethylene metabolic and biosynthetic processes (*p*_Nordic_=1,4e^-3^; *p*_CentralEurope_=9.7e^-4^), jasmonic acid mediated signaling pathways (*p*_SouthWestScandinavia_=1.8e^-2^; *p*_CentralEurope_ =4.7e^-3^; *p*_Scotland_=1.5e^-2^), and intramembrane Golgi-vacuolar transport (*p*_Nordic_=2e^-3^; *p*_SouthWestScandinavia_=4.3e^-3^; *p*_Scotland_=1.8e^-2^). Other relevant processes particular to a specific subpopulation include alkene biosynthesis (*p*_Nordic_ =1,4e^-3^; *p*_CentralEurope_=9.7e^-4^), water transport (*p*_SouthWestScandinavia_ =6.8e^-3^) or secondary shoot formation (*<u>p</u>*_CentralEurope_=3e^-3^).

### Demography and secondary contacts

The expansion of *P. tremula* across the Eurasian continent provides a good scenario to test for changes in population sizes across time. We evaluated population size changes through time using the stairway plot method, which is based on the composite likelihood of the SFS of the species and/or selected subpopulations. We used 50 random samples from our collection to generate the overall behavior of the species, as well as for specific collection sites (Russia and Northern Scandinavia). We observed the strongest reduction in N_e_ around ∼700-800 ka this sharp decrease in N_e_ was observed across all subpopulations, which means it was a bottleneck that affected the entire species and that predated its dispersal in the Eurasian continent. A second bottleneck occurred approximately ∼150-170 ka in the populations from Central Europe and Scandinavia, and a third, mild bottleneck spanned from ∼10-35 ka, a period representing the last glacial era in the Northern hemisphere. These two bottlenecks were not present in the Russian subpopulation and instead, we only observed a second decrease of N_e_ around 60-80 ka in this population (Figure 6).

**Figure 6.**
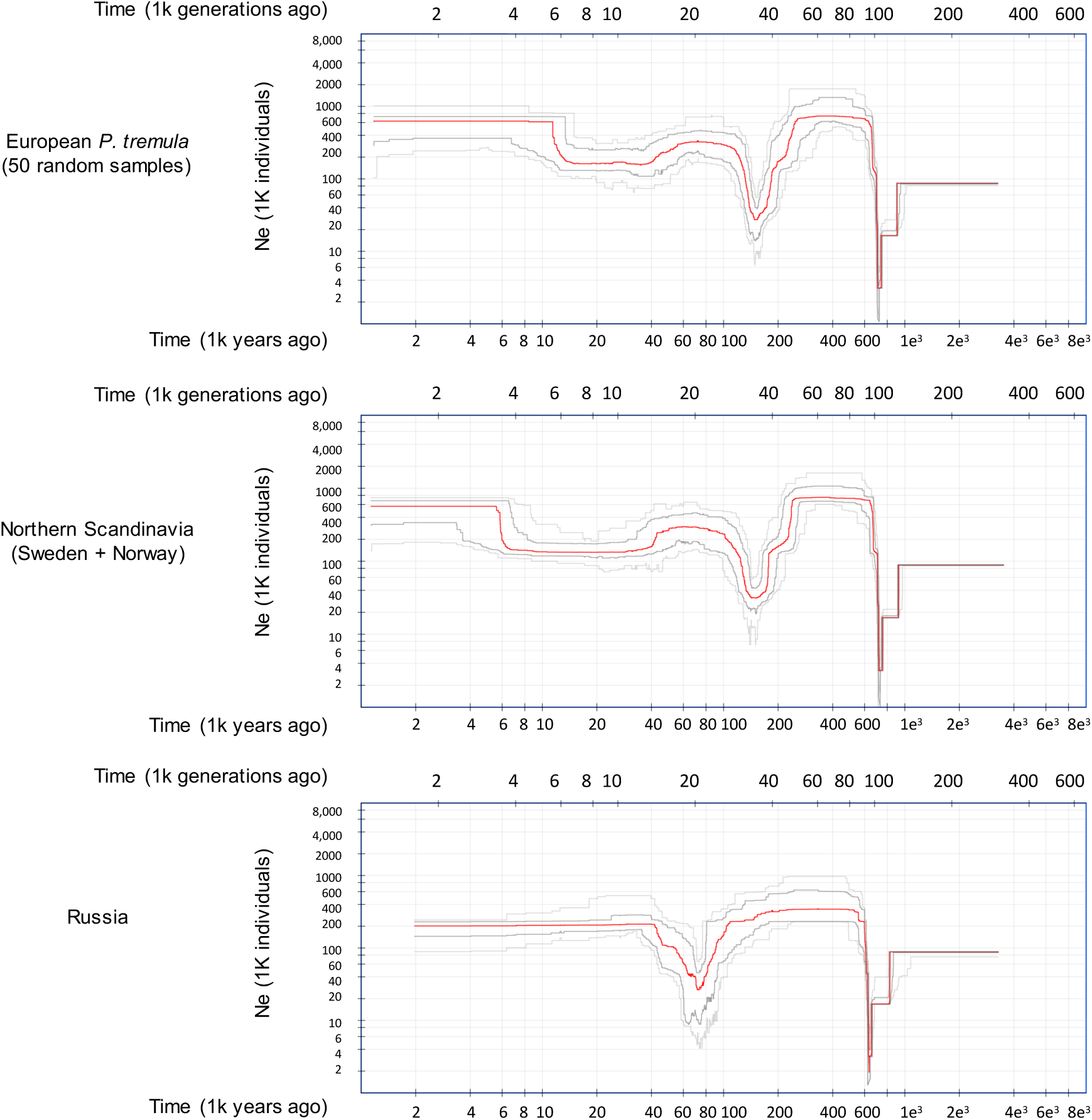
Demographic changes in P*. tremula*.

Given the admixed nature of the Scandinavian individuals, we also evaluated 19 different demographic models including simple models of divergence with and without migration, models with instantaneous size changes, ancient migration or secondary contact, ancient migration plus instantaneous size change, and island models of vicariance and founder events. We used two levels of resolution for this analysis: we ran the 19 models at the chromosome level, using unlinked sites from chromosomes 8 and 10, and also, at a region-targeted level, using the SNPs derived from the segment of chromosome 10 that we hypothesize is the result of adaptive introgression in the Northern Scandinavian population. For these analyses, we considered two populations from our collection, Scandinavia and Russia, in order to identify the most probable site of origin of the selective sweep on chromosome 10. While using all unlinked polymorphic sites on chromosomes 8 and 10 (Figure 7; Additional File1: Figure S10), we consistently observed that the most strongly supported model included divergence in isolation with continuous symmetrical secondary contact and an instantaneous size change (sec_contact_sum_mig_size). Interestingly, at the sweep level, several models were equally supported, with divergence in isolation with continuous asymmetrical secondary contact (with or without instantaneous size change) being the most significant models. Not only was asymmetrical migration detected, but it was stronger from Russia into Scandinavia than in the other direction (Additional File1: Figure S11). After extrapolating the results of these simulations to time units, the secondary contact between Russian and Scandinavian populations was dated to around 30 ka (±3.9e^4^), while the divergence of both lineages occurred 760 ka, consistent with the first bottleneck seen at the species level in the stairway plot analyses.

**Figure 7.**
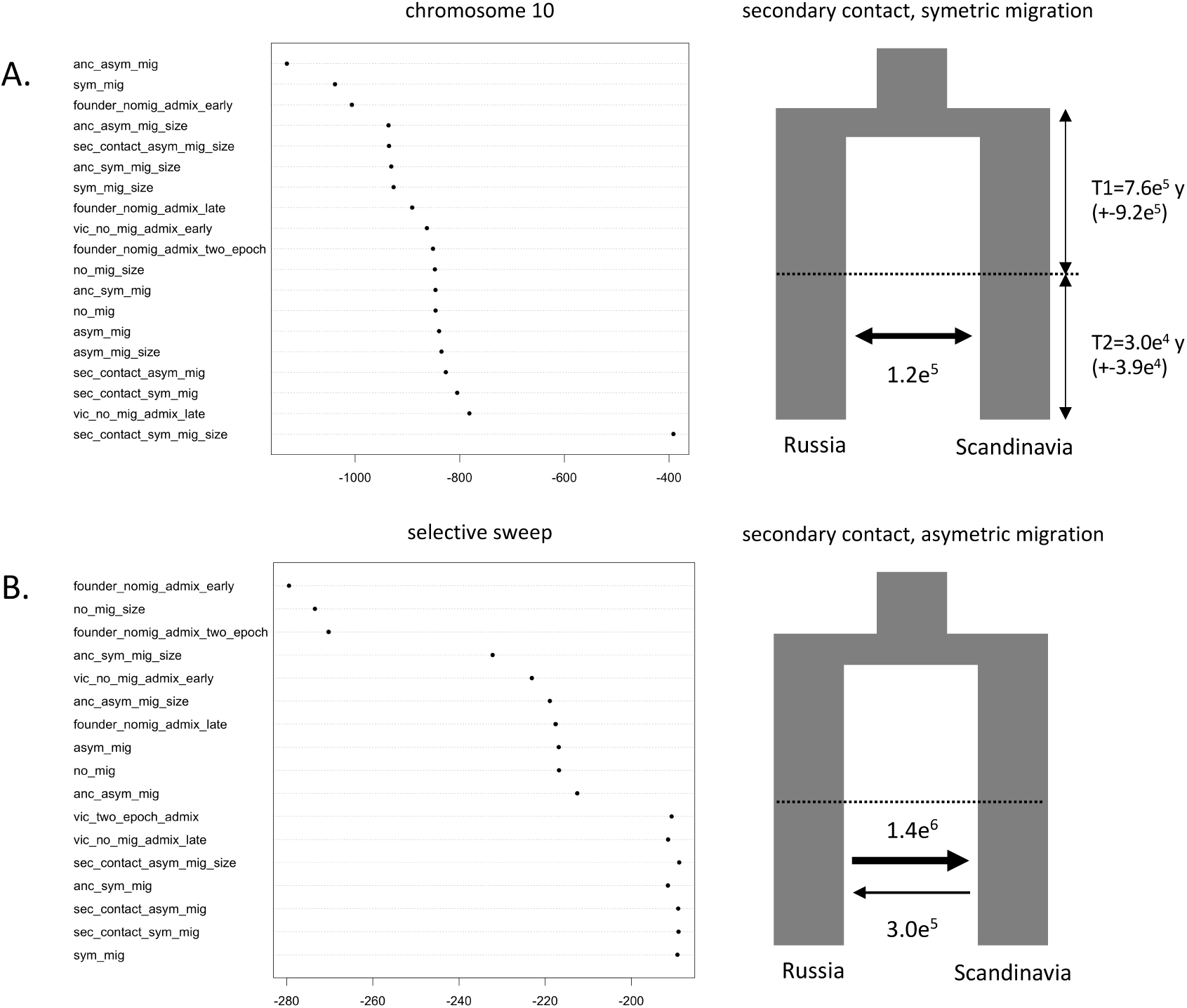
Likelihood of demographic models fitting the SNP data. A. SNPs observed in chromosome 10 are evaluated. B. SNPs comprised within the selective sweep in chromosome 10 are tested.

## Discussion

### Aspen populations: a mosaic of ancestral gene pools

The large number of aspen individuals included in our collection allowed us to dissect the genetic mosaicism of the species across a large geographic scale. Intra-species admixture in *Populus tremula* is not a novel idea, in fact, it has previously been documented between divergent lineages in Europe using data from microsatellite loci [15]. These results, first described in 2010, are, to a certain extent, corroborated by our current analyses but we have also been able to study the hybridization and introgression patterns at much finer scales across the genome of *P. tremula*. The NGSAdmix analysis identified five genetic components that explain the population structure along the East-West geographic axis in Eurasia, corresponding to Eastern Asia (China), Russia, Scandinavia (Sweden and Norway), Eastern European (Latvia) and what we suggest may correspond to Central and Western Europe (Scotland and Southern Norway + Iceland). While Chinese and Russian individuals appear to be partly isolated from Western populations (genome wide F_ST_ values between China and Western Europe ∼0.01, an order of magnitude higher than any other intra-species pairwise comparison), the genomic mosaicism is evident particularly in Scandinavia, where the population can be divided into at least two different groups that follow the latitudinal gradient along the peninsula. Individuals from Northern Scandinavia have a strong Russian ancestral component, whereas those from Southern Scandinavia have strong influence from both Eastern and Central/Western European population. The observations of two independent routes of colonization of Scandinavia, one from the south and another one from the northeast along the ice-free Norwegian Atlantic coast, agree with findings in other organisms, such as brown bears (reviewed by [40]) and humans [41].

de Carvalho and collaborators [15] considered Scottish and Russian subpopulations as proxies for the ancestral populations contributing to the admixed Swedish population. In our analyses we extended this idea by using the Russian, Latvian and Scottish populations in a triple model of hybridization that gave us a higher resolution for identifying admixture routes of the Scandinavian population. In the south of Scandinavia, we could differentiate two groups of individuals: one in Sweden with a clear Latvian component, and one in Norway, with Latvian but also with a Scottish component. This could likely be explained by independent waves of colonization from the south of the peninsula, where Swedish and Norwegian populations were kept partially isolated by the Scandinavian Mountains, or Scandes, that run along the entire peninsula. In line with previous observations, the individuals from the Scottish populations are largely independent from individuals derived from Central Europe. Unfortunately, the lack of samples from Central Europe and the Iberian Peninsula in our collection make it difficult to identify the origin of this population, although de Carvalho et al (2010) identified that trees from Spain constitute a different lineage themselves, possibly due to human-mediated disturbance. Finally, it is clear that the relatively few aspen clones that occur in Iceland are all recent introductions to the island of individuals with a Norwegian origin.

Many very interesting discussions have been carried out in recent years regarding paleographic migration trajectories of trees in Europe [42, 43]. By tracking plant macrofossil and pollen records, it has been suggested that a long-term decline in tree populations occurred around 24 ka followed by a recovery in population sizes approximately 15 ka [7]. The demographic changes that have affected *P. tremula* across the Eurasian continent vary along the East-West axis. The populations from Western Europe have undergone two episodes of bottleneck and subsequent recovery, one around 150 ka and another one during the Middle Pleniglacial period, from which it recovered to its current effective population size between 12-15 ka (Figure 6). On the other hand, the Russian population experienced a single severe bottleneck around 80 ka, from which it recovered before the LGM (Figure 6). The asymmetry in the distributions of tree species along the West to East axis in Europe has been also highlighted by paleobotanical evidence and climate simulations. Western Europe generally lacked tree species north of 46N while higher growing-season warmth in Eastern Europe resulted in increased permafrost thaw and higher water availability so that small populations of boreal trees were able to survive up to approximately 49.8N. The lack of a second bottleneck during the LGM in Russia suggests that the split of the European lineages of aspen predated the glacial maximum and suggests different colonization routes of Russia from southern aspen populations located in far eastern Europe and central Asia.

### Gene flow from the East facilitated adaptation to extreme latitudes

Evidence of inter-species hybridization in poplars has been extensively documented in recent years [14, 44] as well as intra-species gene flow in *P. tremula* [45]. Under these circumstances, we can think of two opposite outcomes of gene flow: it can either homogenize populations, and thus interfere with processes of local adaptation, or it can introduce genetic variation at higher rates than mutation would in the same time frame, providing a source of novel alleles. If any of those foreign alleles are permanently incorporated in a population as a result of gene flow and successive backcrossing, a process defined as introgression, this can increase the fitness of the recipient population and thus, can be referred to as adaptive introgression. Compared to neutral introgression, where alleles can be lost by drift, adaptively introgressed alleles are maintained by selection that can lead to fixation in the recipient population [46]. The role of introgression for species adaptation and evolution has been recognized in animals (reviewed by [47], humans [48], and plants [49, 50]. A recent clear example of adaptive introgression was reported between two species of cypress in the mountainous region of the eastern Qinghai-Tibet Plateau, where loci from *Cupressus gigantea* introgressed into *C. duclouxiana* and thereby facilitated adaptation in *C. duclouxiana* to cooler and drier conditions at higher latitudes and elevations [51].

As several of the populations in our aspen collection are hybrids, we evaluated the occurrence of adaptive introgression events between lineages. One noteworthy genomic region was identified as it displayed clear patterns of introgression between Northern Scandinavia and Russia that were detected using different estimators of introgression (Bd_f_, f_dM_, and local ancestry; Figure 3) and that we could validate using patterns of genetic diversity and differentiation (Additional File1: Figure S6,7). This genomic tract is located on chromosome 10 and comprises ∼500 Kb surrounding two FT homologs that have already been implicated as a key component for seasonal adaptation of phenology in aspens to high latitudes in Sweden [11]. Not only did the genomic tract display a clear signal of positive selection in Sweden, corroborating earlier studies [11], it also showed strong signs of selection in other high-latitude populations, including northern Norway and Russia (iSAFE>0.2; Figure 4).

In order to determine the population of origin of the adaptive allele in this region on chromosome 10, we used demographic modeling to test for different patterns of contact between ancestral populations and directionality of the hypothesized gene flow. First, using a set of unlinked SNPs derived from two independent aspen chromosomes (chr10 and chr8), we observed that Russian and Northern Scandinavian populations have experienced symmetric migration following a secondary contact, making it possible for backcrossing to occur and thereby promoting introgression between the two populations. Second, using information only from the adaptively introgressed region on chromosome 10, we confirmed that, indeed, the secondary contact models are the most strongly supported (together with vicariance and late admixture) and that the migration in this region was stronger from Russia into Scandinavia than in the opposite direction (Additional File1: Figure S10,11). These analyses strongly suggest that regions surrounding the two FT homologs have been adaptively introgressed from the Russian population into Northern Scandinavia in the recent past, thereby facilitating adaptation in *P. tremula* to grow under high latitude conditions.

At the phenotypic level, there is convincing evidence that the *FT*2 allele present in northern Scandinavia displays partial dominance [11]. Trees in the northern part of Scandinavia carrying the northern *FT*2 allele initiate growth cessation about 30 days earlier than trees carrying the southern allele. Trees carrying the northern allele cannot grow in Southern Scandinavia as their critical photoperiod for growth is never reached [52]. This implies that the Nordic allele could have emerged and been kept in the Russian population in a heterozygous form during the last glaciation. The degree of dominance influences the evolutionary dynamics of alleles in diploid populations as the fixation probability of a newly arise (partially) dominant beneficial allele is higher than for a recessive allele in static or slowly changing environments, a process known as Haldane’s sieve [53]. Furthermore, recent modelling suggests that the relationship between dominance and selection coefficients arose as a natural outcome of the functional importance of genes (i.e., degree of connectivity in protein networks) and their optimal expression levels [54]. Given the central role of *FT*2 in regulating the plant phenology signaling pathways and the fine tuning of its expression along plant development, we can speculate that the effect on fitness of the Nordic allele allowed a rapid increase in frequency after the colonization of the Scandinavian peninsula, where it was strongly selected for, allowing a rapid local adaptation to high latitudes. It is worth noting that despite the high recombination rate in *P. tremula* [55], the introgression signal has not been broken and is detectable in a long tract of ∼500 Kb, reinforcing the idea of recent contacts and hybridizations between Nordic populations.

### Other paths to local adaptation

A major genetic process contributing to local adaptation in many populations is that of selective sweeps. Selective sweeps result in the loss of genetic variation in the neighboring regions of a newly adaptive allele or mutation as selection drives this allele to fixation. Selective sweeps generally fall into two categories, hard sweeps and soft sweeps, depending on the origin of the adaptive allele. Hard sweeps are the result of positive selection acting upon a newly arisen beneficial mutation before divergence of lineages, thus the same variation in neighboring haplotypes is driven to high frequencies alongside the adaptive allele. Conversely, a soft sweep occurs from standing variation so that haplotypes surrounding a future beneficial allele have had time to diverge before the onset of positive selection, meaning that the haplotypes that hitchhike together with the beneficial allele may differ between lineages or populations [56, 57].

Our scans for positive selection using the iSAFE algorithm revealed a very strong signal of a selective sweep located on chromosome 10. While we also observed other regions across the aspen genome that seemly have experienced selective sweeps, the calculated iSAFE values in these regions were generally too weak to pass the significance threshold and did not display a clear geographic pattern, which precluded them from being interpreted as selective sweeps involved in conferring local adaptation. A natural conclusion from the lack of non-neutral outliers in our analyses is that local adaptation in aspens is primarily polygenic and/or driven by natural selection acting on standing variation rather than from new mutations that would induce hard sweeps, as is the case for the hard sweep identified on chromosome 10.

The role of standing variation in adaptive evolution has been widely debated. As highlighted by [15], beneficial alleles that are present as standing variation are generally older than new mutations, implying they could have undergone a selective filter during past environmental conditions or in different parts of a species range. These polymorphisms can be maintained for a long time due to balancing selection that persists for many generations and minimizes the effect of drift. When we think of temperate tree species, such as aspens, that have experienced several population contraction and expansion cycles throughout the last millennia, such “pre-selected” standing variation could represent a very useful pool for adaptive alleles that can rapidly be brought together to mediate local adaptation to novel environmental conditions encountered during range expansion. For instance, one climatic consequence for plants during the LGM was water stress due to low CO_2_ concentrations and the presence of permafrost. Even if boreal trees can grow on continuous permafrost, other factors such as soil texture or timing of the spring-summer thaw determine the amount of water available for growth, thus influencing the species distribution [7].

We evaluated the importance of standing variation for local adaptation in aspen by scanning the genome for signals of balancing selection (BS). The combination of significant beta-scores and the increased genetic diversity (π) and pairwise absolute divergence (D_XY_) in several regions across multiple chromosomes, shared by at least two aspen lineages, indicated that BS has maintained ancestral polymorphisms in the species. Indeed, excess diversity around selected loci may be due to the retention of ancestral polymorphisms or due to the accumulation of derived polymorphisms, which happens faster than expected under neutrality. The former is expected if the same allelic lineages have been evolving under BS for a long time, with drift and mutation determining polymorphism within each lineage. The latter would occur if new, more recent instances of BS appear, facilitating the establishment of new neutral polymorphisms [2]; given that our sampling belongs to one single species across the Eurasian continent, we can speculate it is the second scenario which has produced the excess of diversity. Even though gene flow is an important process that could mimic trans-lineage polymorphisms and shared haplotypes between subpopulations, our scans of genomic introgression showed no evidence of such pervasive events, except for the Scandinavian and Siberian lineages.

As reported in other species, the BS regions identified in aspens harbor potentially interesting genes for mediating local adaptation. Of particular relevance was the observation of an enrichment of genes in these genomic regions related to ethylene metabolic and biosynthetic processes. This is of note given that this phytohormone plays a pivotal role in defense responses, plant growth and senescence [58] and, most importantly, in xylem development and the formation of tension wood in aspens [59, 60]. Other functional categories such as jasmonic acid mediated signaling pathways and endo-membrane trafficking suggest differential responses to abiotic stresses that have been favored by the maintenance of advantageous alleles across the species range. In addition, the observation of an enrichment of alkene biosynthesis related genes is intriguing, as a significant association between the presence of alkenes in cuticular waxes and tree growth and resistance to leaf spot have been reported in *P. trichocarpa* [61]. Thus, we suggest that standing variation and balancing selection have shaped the capacity of aspen populations to adapt to a wide range of environmental conditions during the post-glacial colonization of Europe.

## Conclusion

Our large panel of re-sequenced aspen individuals allowed us to unravel key genomic aspects behind local adaptation across the Eurasian continent. It is clear from our results that intra-species hybridization has played a major role homogenizing the genomic background of the species and has promoted the movement of adaptive alleles between populations. Of major relevance is the observation of a recent adaptive introgression event between Nordic populations around the *Flowering locus T*, that has facilitated the survival of aspens in high latitudes. This event is however the only such event that can be detected in the species, showing that the emergence of advantageous alleles and their propagation is rather rare. Standing variation and its conservation through balancing selection across lineages seems to be a more efficient way of keeping advantageous alleles and maintaining high levels of diversity. This combination of evolutionary scenarios suggests that aspens may have the capacity to adapt rapidly to new challenging environments, and this augurs well for the survival of the species under a range of possible future climate conditions.

## Acknowledgments

The SwAsp common garden was maintained by Skogforsk. The Scottish clone collection was commissioned by Forest Research and NatureScot, and planted and maintained by Forest Research Technical Services Unit, with funding from the Forestry Commission. We thank Peter Livingston and Eadha Enterprises for the samples from the EADHA clone collection. We are grateful to the Bioinformatics facility at Umeå Plant Science Centre for curation of sequencing data. SJ, KMR and NRS are supported by the Trees for the Future (T4F) project. PKI, SJ and KMR are supported by the Knut and Alince Wallenberg Foundation (KAW). The research was also supported by grants from the Swedish Research Council (VR) to PKI and to SJ, and by Formas to SJ. Part of the data generated for the study was supported by Science for Life Laboratory and the National Genomics Infrastructure (NGI, as a SciLifeLab National project) which provided access to massive parallel sequencing. All analyses were performed on resources provided by the Swedish National Infrastructure for Computing (SNIC) at Uppsala Multidisciplinary Centre for Advanced Computational Science (UPPMAX) under compute projects 2017/7-219, 2018/3-552, 2019/3-597, 2020/5-621 and storage projects sllstore2017050, sllstore2017059.

## Data accessibility

All raw read data have been uploaded to NCBI/ENA and can be found under Bioproject IDs PRJNA297202 and PRJEB42846.

## Author contributions

PKI, NRS, KR, SJ and MR-A planned and designed the research. MR-A, JW and PKI performed experiments and analyzed data. SS, AF, JC, MESB, CL, DR, KR and NRS provided biological material and performed field work. MR-A and PKI wrote the manuscript with input from NRS and KR. All authors read and approved of the final version of the manuscript.

