## Supplementary Materials for "Adaptive introgression and standing genetic variation, two facilitators of adaptation to high latitudes in European aspen (*Populus tremula* L.)"

**Additional File 1.** Supplementary Figures.

Supplementary Fig. S1. SwAsp population structure before (A) and after (B) batch removal. PCAs were performed on pruned SNPs (LD=0.2; MAF=0.05); in B, SNPs contributing more than expected according to our uniformity hypothesis were removed [T=((C*eig1)+(C*eig2))/(eig1+eig2)]. In red, individuals sequenced by Illumina NextSeq; in blue, individuals sequenced by Illumina HiSeq2000.

Supplementary Fig. S2. SwAsp population structure after batch removal. PCAs were performed on pruned SNPs (LD=0.2; MAF=0.05) and applying different p-value filters to the contribution of the variants to PC1 and PC2. In red, individuals sequenced by Illumina NextSeq; in blue, individuals sequenced by Illumina HiSeq2000.

Supplementary Fig. S3. Demographic models tested between Northern Scandinavia and Russia.

Supplementary Fig. S4. Likelihood of the tested model after each round of optimization.

Supplementary Fig. S5. Effect of batch removal in the iSAFE analysis. Manhattan plots of iSAFE values contrasting the Northern and Southern Swedish populations in a case/control configuration, before (left panel) and after (right panel) batch removal (BR).

Supplementary Fig. S6. Pairwise $D_{XY}$ at genome-wide, selective sweep and BS regions.

Supplementary Fig. S7. Pairwise $F_{ST}$ at genome-wide, selective sweep and BS regions.

Supplementary Fig. S8. Allele dosage along chromosome 10 estimated in five independent ELAI replicates. In green, the dosage from the ancestral Russian population; in brown from the Latvian population.

Supplementary Fig. S9. iSAFE pattern across populations.

Supplementary Fig. S10. Likelihood of the demographic models. Simulations using all unlinked sites on chromosomes 8 and 10.

Supplementary Fig. S11. Likelihood of the demographic models. Simulations using sites on the selective sweep on chromosome10, changing the directionality of the migration.

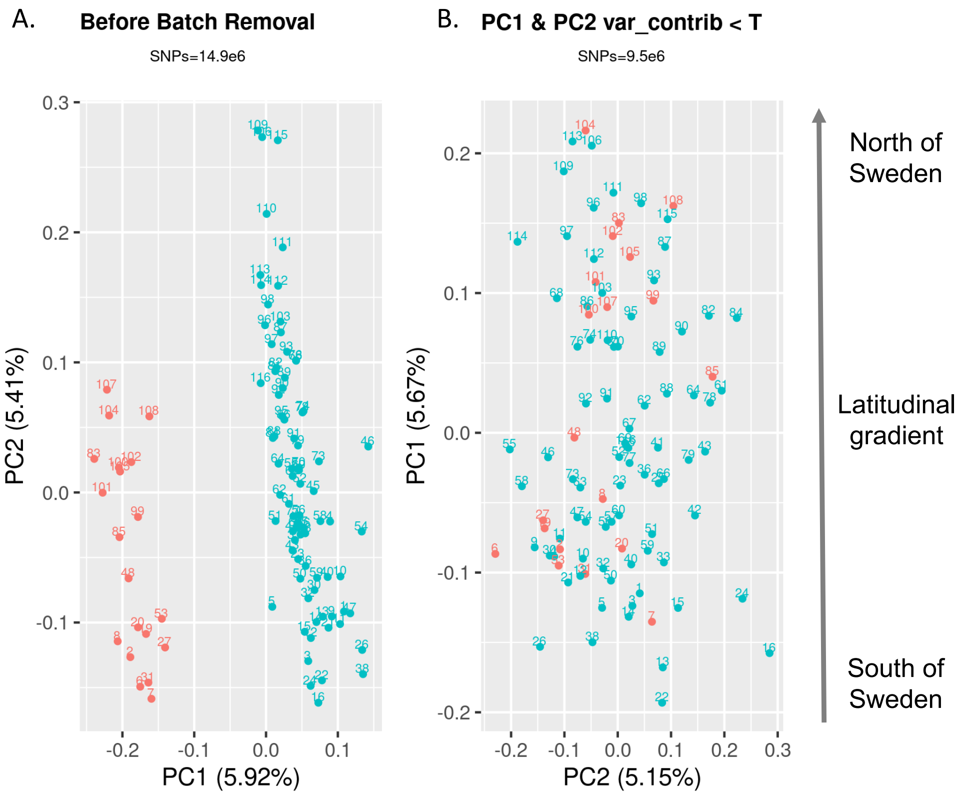


Supplementary Fig. S1

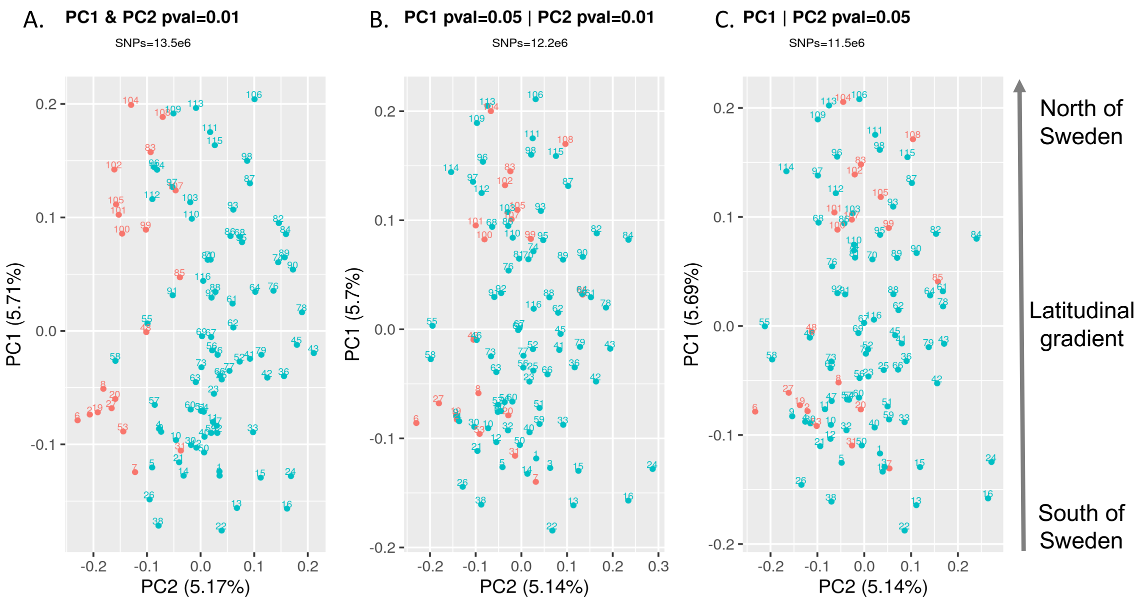


Supplementary Fig. S2

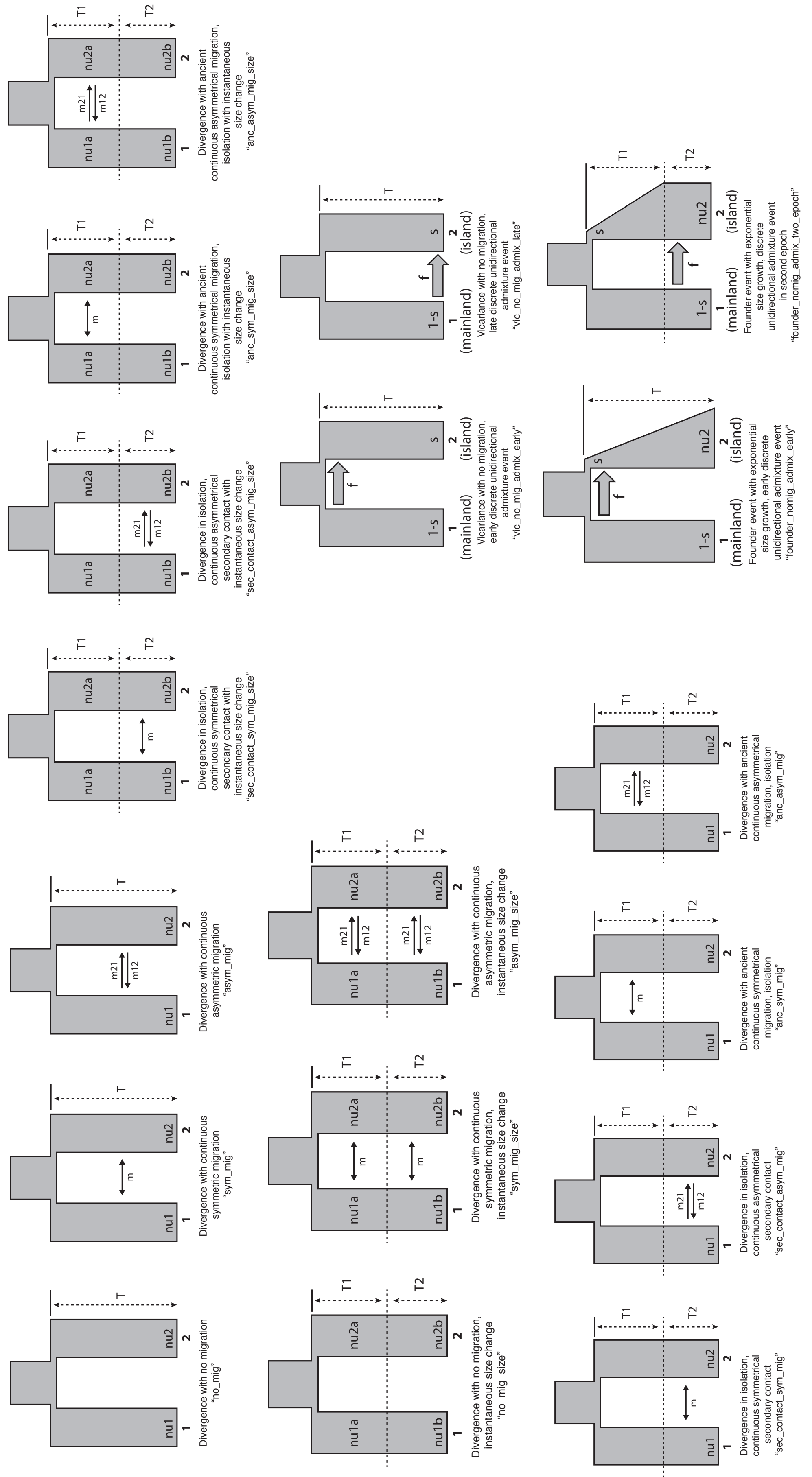


Supplementary Fig. S3

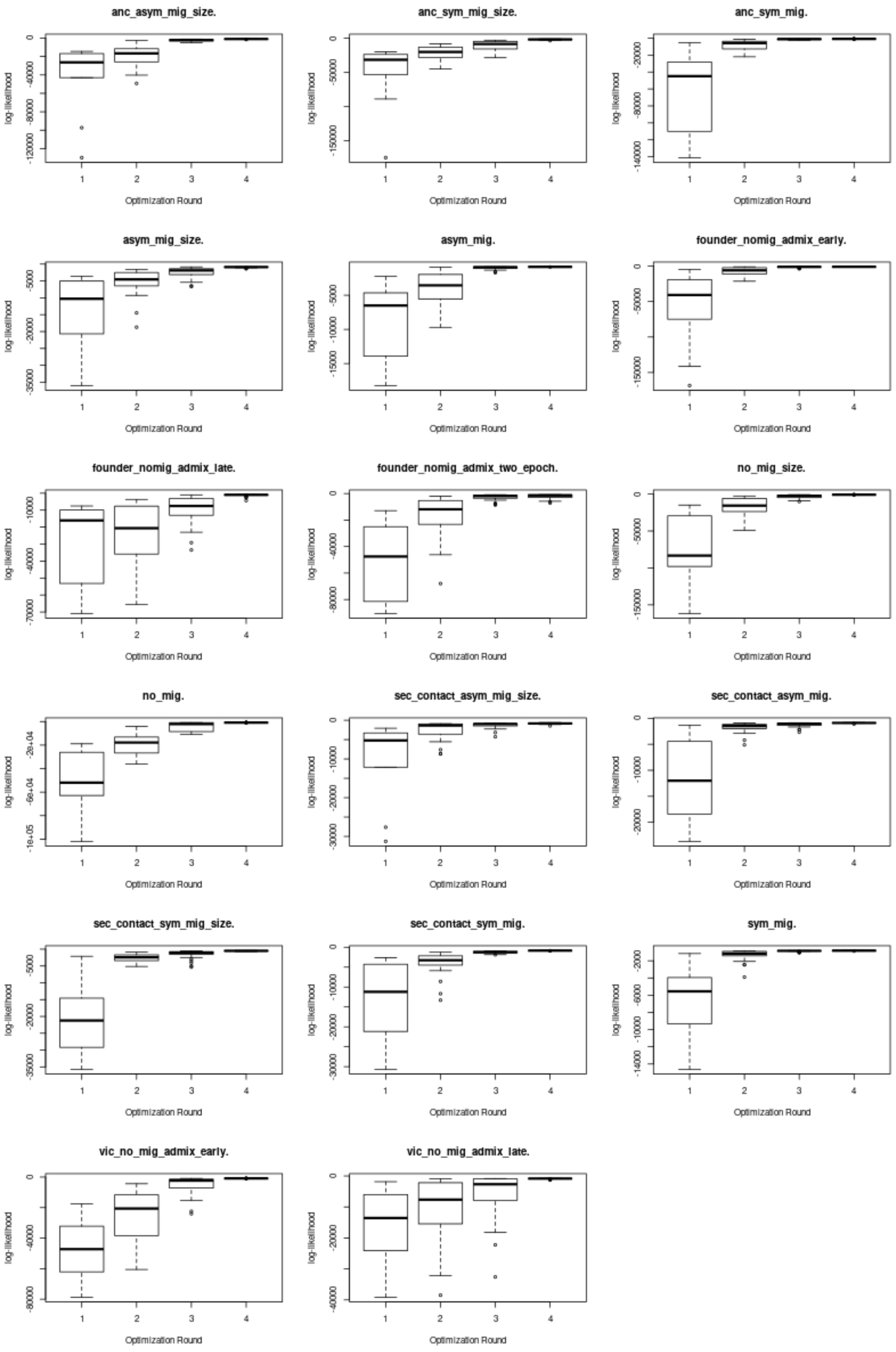


Supplementary Fig. S4

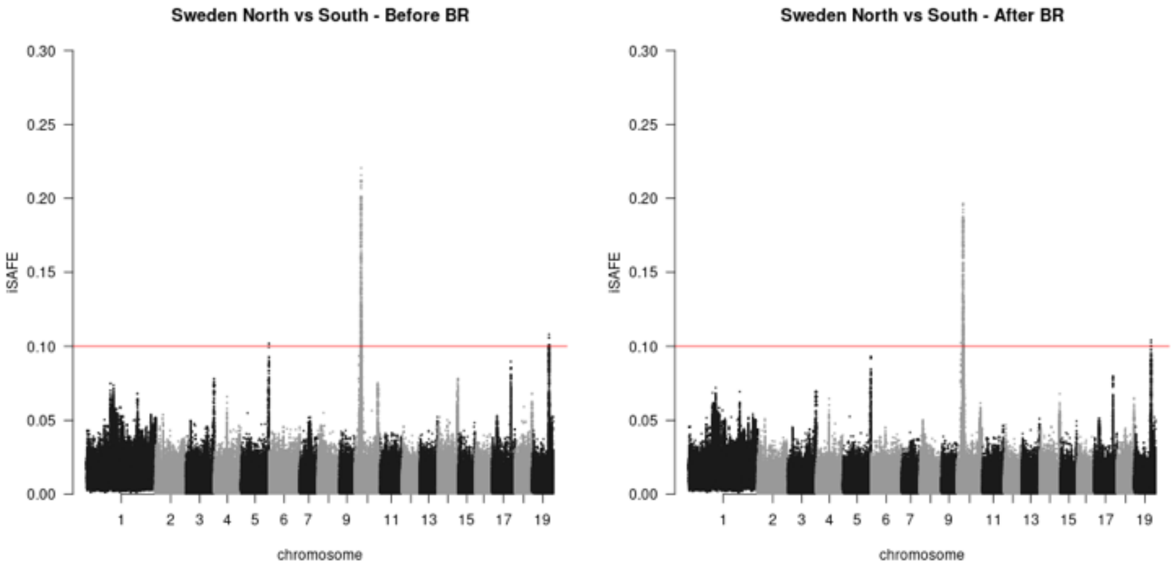


Supplementary Fig. S5

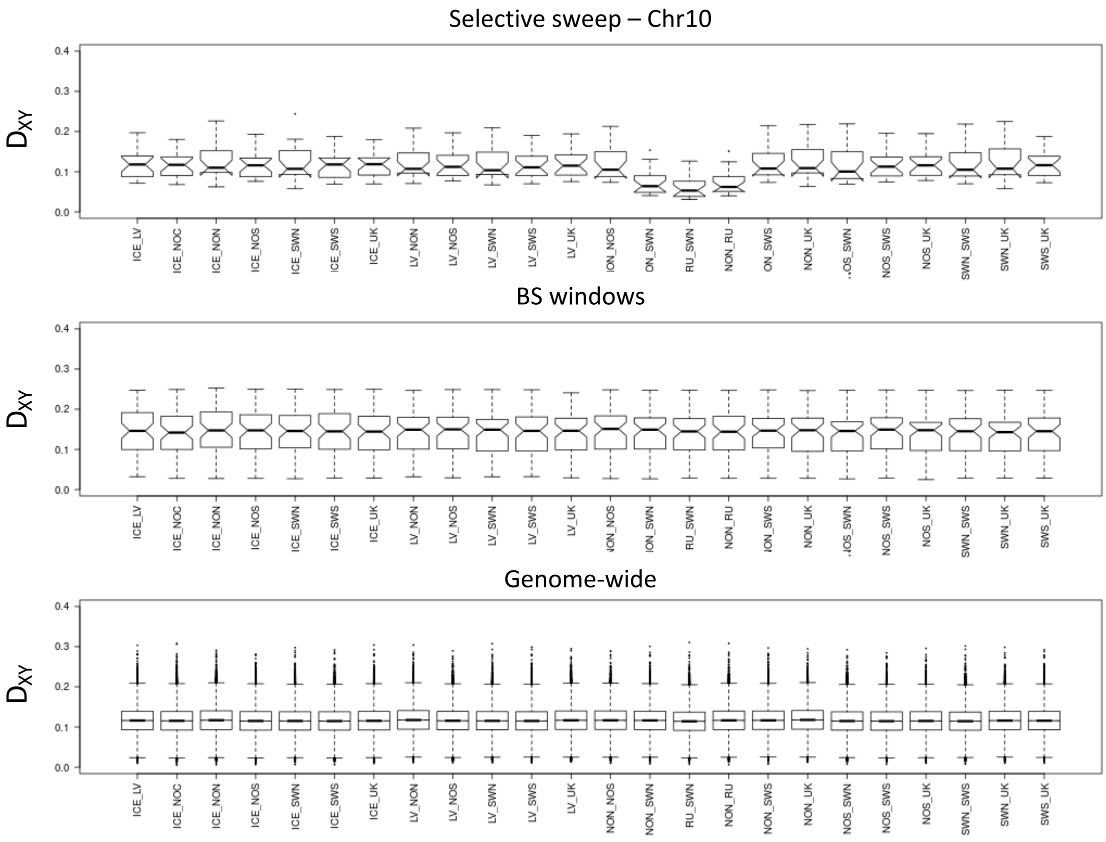


Supplementary Fig. S6

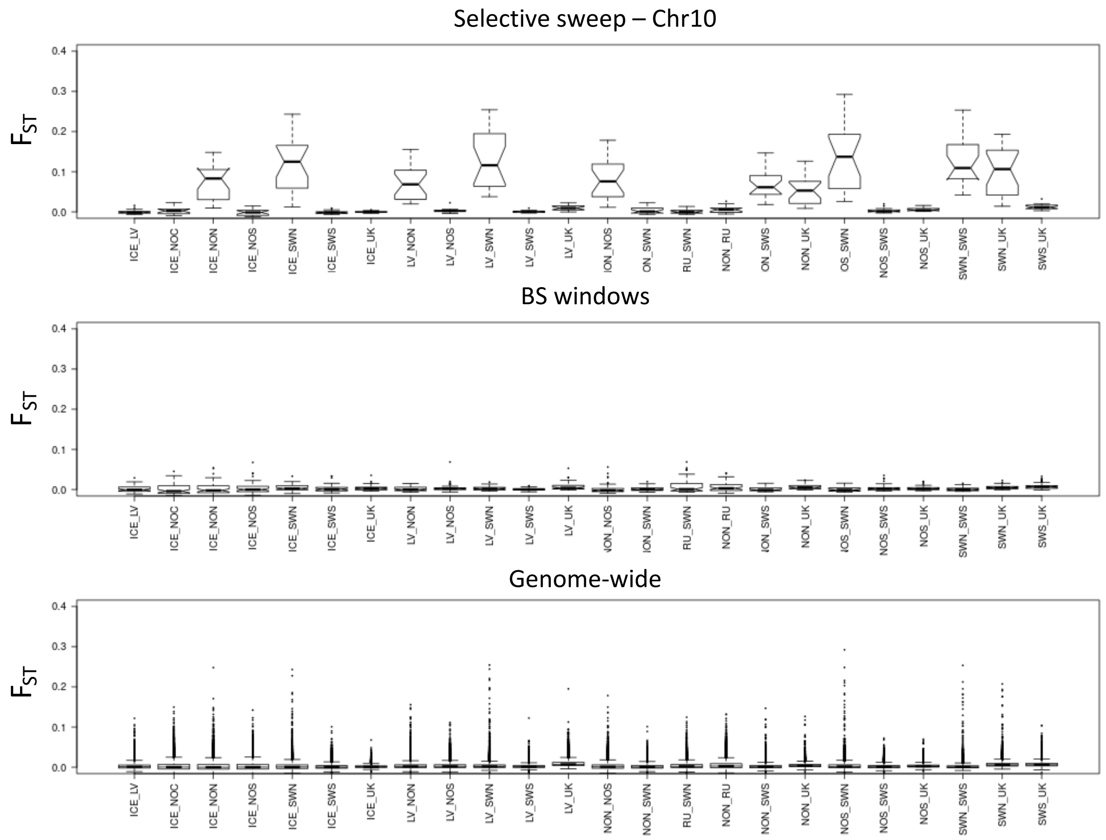


Supplementary Fig. S7

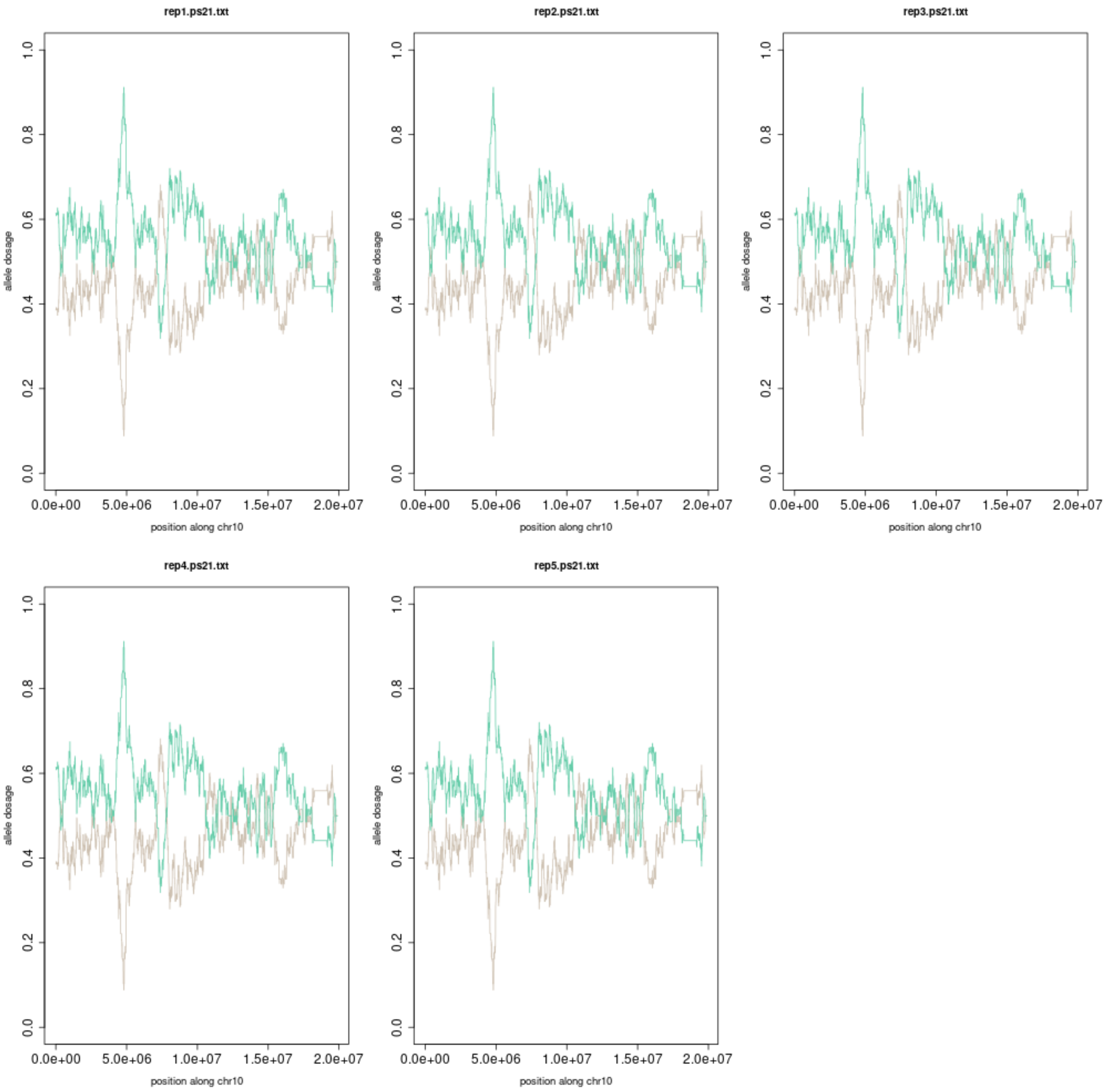


Supplementary Fig. S8

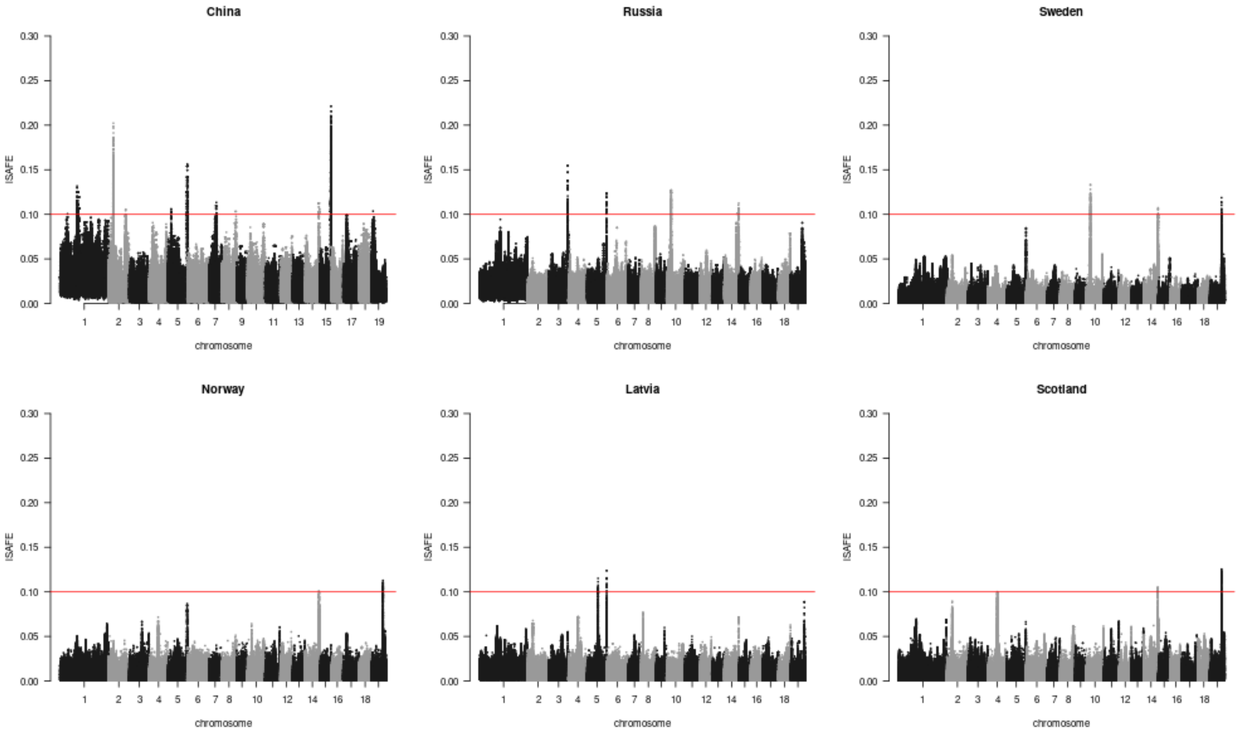


Supplementary Fig. S9

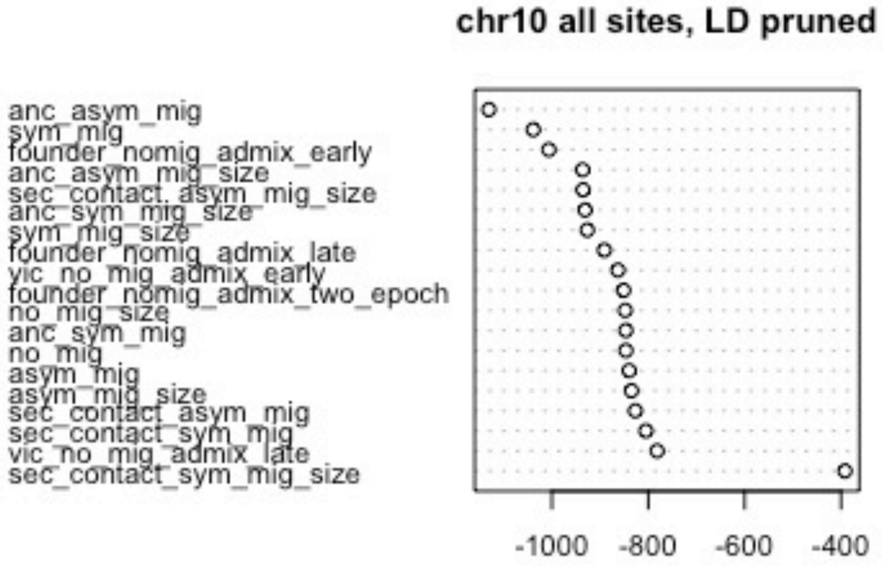


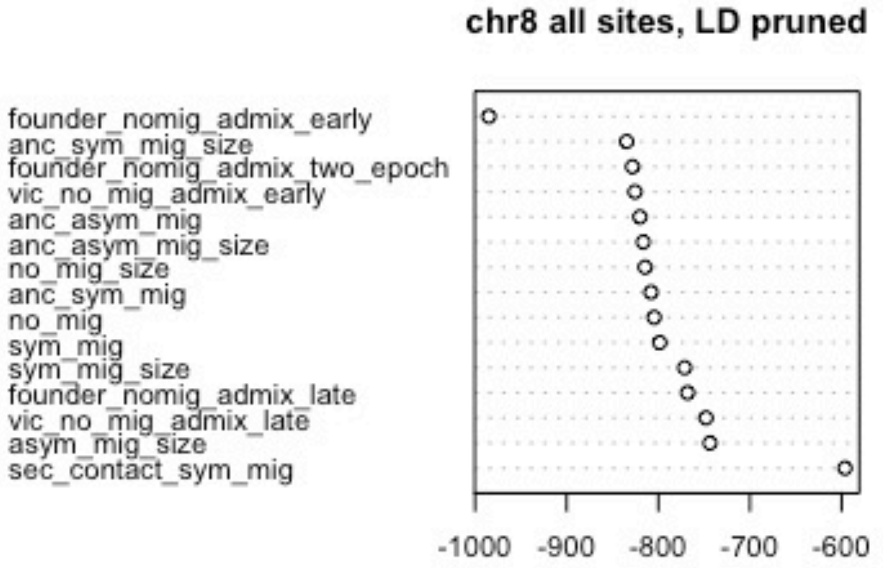


Supplementary Fig. S10

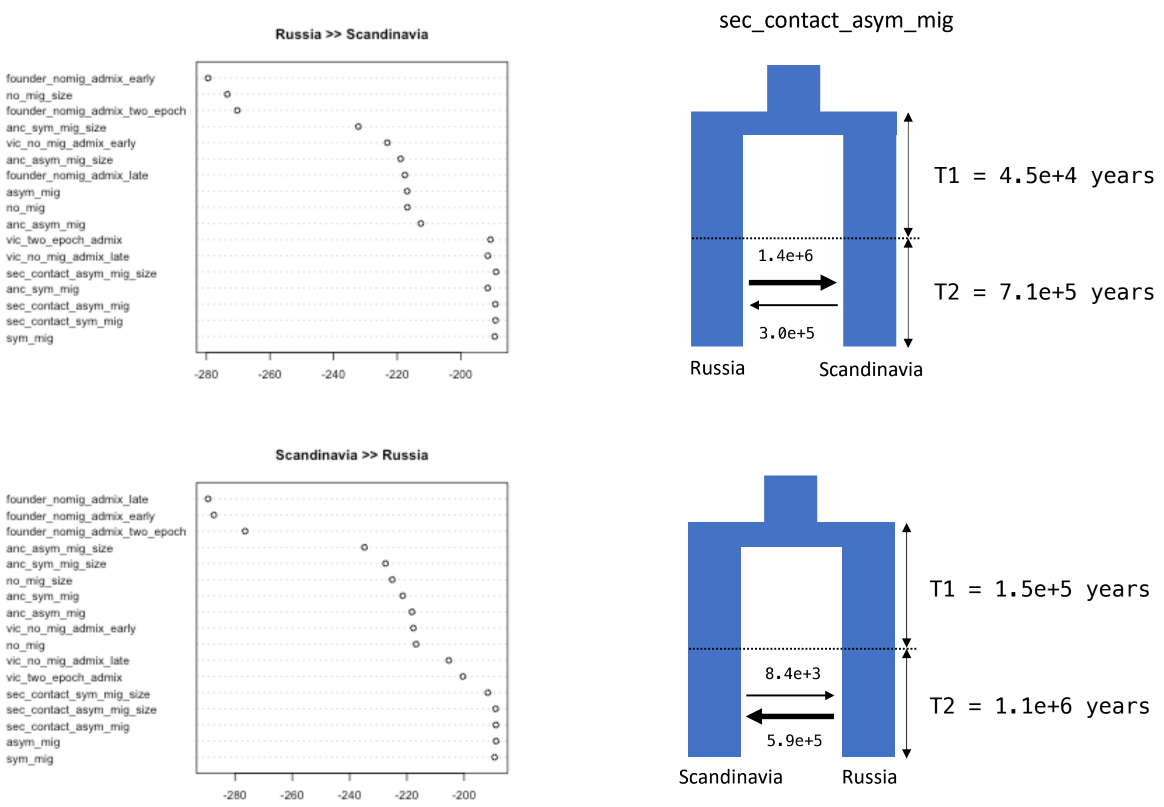


Supplementary Fig. S11
